# Rust fungi secretomes contain structurally diverse effector families including cysteine-rich metal-binding proteins

**DOI:** 10.64898/2026.06.11.731755

**Authors:** Megan A. Outram, Zhao Li, Michael Kuiper, Simon J. Williams, Melania Figueroa, Peter N. Dodds, Jana Sperschneider

## Abstract

Rust fungi are significant threats to global food security, causing substantial damage to crops through their ability to adapt and evolve new strains that overcome resistance. These obligate biotrophs infect host plants by secreting effector proteins that manipulate host physiology to promote infection and colonisation. We used AlphaFold2 to investigate structural conservation among effector proteins for the secretomes of *Melampsora lini* and four *Puccinia* species. AlphaFold2 yielded high-confidence predictions for 45.7% of the 27,090 secreted proteins, while 19% were poor quality. Comparative analysis revealed extensive structural diversity across the rust secretomes, with all thirteen known rust Avr proteins belonging to different clusters apart from AvrSr13 and AvrSr33. Nevertheless, there were still numerous large clusters of structurally-related proteins, including 59 clusters with over 50 members each, three of which contained known Avr proteins. Of the major structural families defined in other fungi, the rust species studied here only contained FOLD and ToxA-like families. Structural analysis of cysteine-rich proteins revealed over a thousand effector candidates featuring zinc-binding sites, with approximately 75% predicted to be cytoplasmic effectors. In contrast, cysteine-rich apoplastic effector candidates were characterized by a high frequency of disulfide bonds. One family of predicted metal-binding proteins was greatly expanded in *P. graminis* f. sp. *tritici* and includes AvrSr13 and AvrSr33. We confirmed that purified AvrSr13 and AvrSr22 proteins bind to zinc *in vitro* using biochemical assays. Taken together, structural modeling provides new avenues to study sequence-unrelated effectors and highlights the high degree of diversity in the effector repertoires of rust species.

## Introduction

Rust fungi pose a significant threat to cereal crop production and global food security (Duplessis et al., 2021). Like other phytopathogens, rust fungi secrete effector proteins that act either in the apoplast or are translocated into host cells, where they function to facilitate infection and host colonisation (Figueroa et al., 2020). However, some effectors, known as avirulence (Avr) proteins, are recognised by specific immunity receptors in plants, triggering resistance to the pathogen (Bentham et al., 2020; Dodds et al., 2024). As such, understanding the effector repertoire of a pathogen is crucial for informing and developing effective disease resistance strategies. Because effectors are under strong evolutionary pressures to evade detection whilst maintaining or evolving new virulence traits, they typically show high sequence diversity, limited conservation of functional motifs, and little homology to proteins with known functions (Thordal-Christensen et al., 2018). This makes the identification and functional characterisation of effectors based solely on sequence data particularly challenging.

Experimental structure determination of effectors has significantly enhanced our understanding of their function and recognition by host immune receptors (Mukhi et al., 2020; Outram et al., 2022). In oomycetes, structural conservation has been observed in a large subset of RxLR effectors, which contain one or more structurally similar WY domains (Boutemy et al., 2011; Chou et al., 2011). Several structurally related effector families have also been found to include sequence-unrelated effectors from various fungal taxa. These include the MAX (*Magnaporthe oryzae* Avr effectors and ToxB) (de Guillen et al., 2015; Lahfa et al., 2024; Le Naour—Vernet et al., 2023), RALPH (RNAse-Like Proteins associated with Haustoria) (Pennington et al., 2019; Spanu, 2017), ToxA-like (Sarma et al., 2005; Wang et al., 2007), LARS (*Leptosphaeria* Avirulence and Suppressing) (Lazar et al., 2022), and FOLD (*Fusarium oxysporum* f. sp. *lycopersici* dual-domain) (Yu et al., 2024) families. The ToxA-like family occurs in a large number of diverse fungal pathogens including ascomycetes and basidiomycetes (e.g. *Pyrenophora tritici repentis, F. oxysporum* f. sp. *lycopersici, Melampsora lini*) (Di et al., 2017; Sarma et al., 2005; Wang et al., 2007). Other effector families have narrower distributions among pathogen species. For example, MAX effectors are identified in only a few ascomycetes but are massively expanded in the rice blast pathogen *Pyricularia oryzae*, where they constitute 5-10% of the secretome and approximately 50% of the known Avr proteins (de Guillen et al., 2015; Seong & Krasileva, 2021). Similarly, RALPH effectors occur as a large family in powdery mildew fungi (*Blumeria* spp.,) and include almost all of the identified mildew Avr proteins (Bauer et al., 2021; Pedersen et al., 2012; Pennington et al., 2019; Spanu, 2017).

The emergence of highly accurate *ab initio* protein structural predictors, such as AlphaFold2 (Jumper et al., 2021), RoseTTaFold (Baek et al., 2021), and trRosetta (Du et al., 2021), has enabled large-scale structural studies and comparison of pathogen effectors. For example, Seong and Krasileva (2021) used trRosetta to model secreted proteins from *Pyricularia oryzae* and found additional members of the MAX effector family as well as numerous other conserved structural families. Lahfa *et al*. (2024) used a combination of AlphaFold2 and experimental structure determination to identify 20 subgroups of the MAX family with variations on the canonical fold, including domain extensions and duplications. These subgroups may reflect extended functional diversity in virulence and host targets previously not captured without large-scale analyses. Two studies identified the presence of FOLD effectors across several fungal genera that exclusively interact with plants, including in the arbuscular mycorrhizal symbiotic fungus *Rhizophagus irregularis* (Teulet et al., 2023; Yu et al., 2024). More extensive cross-species comparisons have identified numerous additional families of sequence-unrelated but structurally similar effectors that are present across diverse fungal pathogens. For instance, Seong and Krasileva (2023) found numerous effector families, including the MAX, RALPH and ToxA-like groups, that were distributed across distantly related species, while other families exhibited species-specific expansions, such as the AvrSr50 and AvrSr35-like families in *P. graminis*. Derbyshire and Raffaele (2023) investigated candidate orphan effectors (containing a signal peptide but lacking a functional domain) across 20 fungal species from the Pezizomycotina subphylum and found that most were contained in 62 structurally conserved families.

In rust fungi, seven Avr effectors have experimentally resolved structures: AvrP, AvrL567, AvrM, and AvrM14 from *M. lini*, and AvrSr27, AvrSr35, and AvrSr50 from *Puccinia graminis* f. sp. *tritici* (Förderer et al., 2022; McCombe et al., 2023; Ortiz et al., 2022; Outram et al., 2024; Ve et al., 2013; Zhang et al., 2018; Zhao et al., 2022). Among these, only AvrL567 shares structural similarity with a known effector family (ToxA-like), while the remaining effectors are structurally distinct. Notably, AvrP and AvrSr27 are cysteine-rich cytoplasmic effectors where the cysteine residues are involved in zinc coordination (Ortiz et al., 2022; Outram et al., 2024; Zhang et al., 2018). Similarly, the *Pyricularia orzyae* AvrPii cytoplasmic effector also binds zinc via cysteine coordination (De la Concepcion et al., 2022). The discovery of zinc coordination in these structures raises the question of whether this feature is a widespread characteristic among cysteine-rich cytoplasmic effectors. This would contrast with the situation for cysteine-rich effectors that function in the apoplast, in which the cysteine residues have been found to be involved in disulfide bridges that may enhance stability in the apoplastic environment (Outram et al., 2021; Rocafort et al., 2020; Yu et al., 2024).

Here, we report AlphaFold2 structure predictions of secreted proteins from five rust fungi, including four cereal-infecting *Puccinia* species and the flax-infecting species *M. lini*. The selected *Puccinia* species include three major pathogens of wheat; *P. graminis* f. sp. *tritici* (causing stem rust disease), *P. triticina* (leaf rust disease), *P. striiformis f. sp. tritici* (stripe rust disease) and one pathogen of oat, *P. coronata* f. sp. *avenae* (crown rust disease). We identified structural families that are conserved within and/or across the rust species, including several expanded families containing known Avr proteins. Using an *in silico* approach we identified expanded families of putative zinc-binding proteins in the rust secretomes and experimentally validated zinc binding for two stem rust effectors, AvrSr13 and AvrSr22. Collectively, our findings provide new insights into the structural diversity of rust effectors, offering a valuable resource for understanding rust infection strategies and host recognition.

## Materials and methods

### Gene annotations and secretome prediction

Secretomes were extracted from genome annotations using SignalPv4.1 (-t euk -u 0.34 -U 0.34) (Petersen et al., 2011) to identify proteins containing a signal peptide. Proteins predicted by TMHMM v2.0 (Krogh et al., 2001) to include one or more transmembrane domains after the first 60 amino acids were excluded. The published genome annotations for *P. striiformis* f. sp. *tritici* strain Pst-104E (Schwessinger et al., 2018), *P. triticina* isolae Pt76 (Duan et al., 2022; Sperschneider et al., 2023) and *M. lini* isolate CH5 (Sperschneider et al., 2025) were used. For *P. graminis* f. sp. *tritici* and *P. coronata* f. sp. *avenae*, the Pgt21-0 (Li et al., 2019) and Pca203 (Henningsen et al., 2022) genomes were re-annotated using the pipeline described for *P. triticina* Pt76 (Sperschneider et al., 2023). EffectorP 3.0 (Sperschneider & Dodds, 2021) was used to predict apoplastic and cytoplasmic effectors in the rust secretomes. For proteins with a dual-localisation prediction, the highest probability from the EffectorP prediction was selected to define localisation. InterProScan (Jones et al., 2014) was used to detect potential PFAM domains.

### Protein structural modelling

Structural models were generated with AlphaFold2 v2.3.1 (Jumper et al., 2021) using the full databases (uniref90, uniref30, MGnify, BFD; as downloaded from https://github.com/google-deepmind/alphafold) to generate the multiple sequence alignments. All structure templates from the protein databank (PDB) downloaded on July 20, 2021, were used for structural modelling. For each of the proteins, we selected the best ranked model for the downstream clustering and structural analysis. To assess the overall confidence of each prediction we calculated the average pLDDT (predicted local distance difference test) score for each protein. Structures were visualised using pymol (https://github.com/schrodinger/pymol-open-source) and coloured using the pymol-color-alphafold extension (code sourced from https://github.com/cbalbin-bio/pymol-color-alphafold).

### Comparison of protein structures and similarity clustering

The ‘createdb’ module of Foldseek v10 (van Kempen et al., 2023) was used to construct a local target database of all the structure predictions of the rust secretomes, along with 58 experimentally determined effector protein structures from various fungal and oomycete pathogens (Table S1). We also included the individual N- and C-terminal domains of AvrSr27 and Avr1/SIX4 as these proteins contain two subdomains (Outram et al., 2024; Yu et al., 2024). Clustering of the rust secretome structures and experimentally determined effector protein structures was conducted using the easy-cluster module of Foldseek (-s 7.5 --tmalign-fast 0 --cov-mode 0 -c 0.5 --alignment-type 1 --tmscore-threshold 0.55).

### *In silico* identification of zinc-binding sites and disulfide bonds

To identify putative zinc-binding sites or disulfide bonds in predicted protein structures, we employed the analyze_AlphaFold2_metal_clusters_for_release Software (https://github.com/Elcock-Lab/Metalloproteome) developed by Wehrspan et al. (2022). Briefly, the algorithm screens protein structures for arrangements of cysteine (C) and histidine (H) residues fitting geometric criteria consistent with either disulfide bonds or with six variations of zinc-binding sites (CHHH, CCHH, CCCH, HHH, CCCC, and HHHH). Putative disulfide bonds were called if two cysteine residues within searched regions were within 2.5 Å of one another. The following parameters were used: thrregionDistanceMinimum (0.0), regionDistanceMaximum(8.0), stericClashCutoff (2.0), disulfideCutoff (2.5), addBbnAtomCutoff (0.0), ligandLigandClashCutoff (2.5). Hydrogen bonds were removed from all model files prior to *in silico* screening using a python script that removed all atoms where element=H.

### Protein expression and purification

The coding sequences of AvrSr13 (residues 21-138), AvrSr22a (residues 24-97), AvrSr22b (residues 24-97) and AvrSr22a^C-trunc^ (residues 24-74) (Arndell et al., 2024) were codon optimised for expression in *E. coli* and synthesised as double stranded DNA gene blocks (Table S2) and subsequently cloned into a modified pOPIN expression vector by Golden Gate cloning (Bentham et al., 2021; Yu et al., 2021). The expression constructs contain an N-terminal 6xHis-tag followed by a 3C protease cleavage site for protein isolation and purification. The pOPIN expression vectors with inserted genes were transformed into BL21 (DE3) competent *E. coli* (NEB, 2527I). Overnight cultures of transformed BL21 (DE3) cells were used to inoculate Terrific Broth media supplemented with trace metals solution (1000X trace metals solution is 50 mM FeCl3, 20 mM CaCl2, 10 mM MnCl2, 10 mM ZnSO4, 2 mM CoCl2, 2 mM CuCl2, and 2 mM NiCI2) (Studier, 2005). Cells were grown at 37°C with shaking at 220 RPM until the optical density at 600 nm reached 0.6-0.8. Cultures were then induced with 250 µM isopropyl-1-thio-β-d-galactopyranoside (IPTG) and the temperature was reduced to 18°C for a further 16 hours culture before the bacteria cells were harvested by centrifugation (4,000 g for 30 mins at 4°C). Cells were resuspended in a lysis buffer containing 50 mM HEPES pH 8.0 for AvrSr13 and pH 7.0 for AvrSr22, 300 mM NaCl, 10% glycerol, 1 mM phenylmethanesulfonyl fluoride (PMSF) and 1 mM dithiothreitol (DTT). Cells were lysed on ice using sonication (5min, 40% amplitude, 10 s on, 20 s off), followed by centrifugation at 18,000 g for 30 min at 4°C. To the clarified lysate, 20 mM imidazole was added before the lysate was applied to a 5 mL HisTrap FF crude column (GE Healthcare). The column was washed with 20 column volumes of wash buffer (50 mM HEPES, pH adjusted depending on the protein, 300 mM NaCl, and 30 mM imidazole) to remove non-bound proteins. The target protein was eluted by a linear gradient from 30 mM to 250 mM imidazole over 10 column volumes. Elution fractions were visualised for protein quantity and purity using Coomassie-stained SDS-PAGE. Fractions containing the protein of interest were buffer exchanged via dialysis in 20 mM HEPES, pH 8 (AvrSr13) or 7 (AvrSr22) and 300 mM NaCl, using 3.5 kDa MWCO SnakeSkin Dialysis tubing. The N-terminal 6xHis-tag was cleaved with Human Rhinovirus 3C protease at 4ºC overnight. The cleaved samples were concentrated by using an Amicon centrifugal concentrator (Merck) with 3kDa MWCO for AvrSr22a^C-trunc^ and 10 kDa MWCO for AvrSr13, AvrSr22a and AvrSr22b. Samples were then further purified using size exclusion chromatography (SEC) on a Superdex 75 16/600 column (GE Healthcare) equilibrated with 20 mM HEPES, pH-adjusted depending on the protein, 150 mM NaCl and 1 mM DTT. Protein concentration was determined by measuring the absorbance at 280 nm and incorporating the extinction coefficient as calculated by the Expasy ProtParam tool (http://ca.expasy.org/cgi-bin/protparam).

### 4-(2-pyridylazo) resorcinol (PAR) assay

PAR assays were carried out as described previously (Ortiz et al., 2022; Outram et al., 2024). Proteins were denatured by boiling at 95°C with 1% SDS for 10 minutes to release any metal ions that could be coordinated by the proteins and then mixed with the PAR solution to a final concentration of 50 μM. An equal volume of buffer was used as blank and the AvrSr50-C protein, which does not bind metals (Ortiz et al., 2022; Outram et al., 2024), was used as negative control. The absorbance of the samples was measured at wavelengths between 350 and 650 nm using a Spectromax Quickdrop (Molecular Devices).

### Metal ion characterisation and quantification via Inductively Coupled Plasma Mass Spectrometry (ICP-MS)

To identify the bound metal ions in purified proteins we performed ICP-MS. Before analysis, proteins were dialysed against one litre of SEC buffer (20 mM HEPES, pH adjusted according to the requirement of each protein, with 150 mM NaCl). Dialysed proteins were adjusted to a concentration of 5 μM and digested with 2% HNO3 (analytical grade) overnight at room temperature followed by centrifugation at 16,000 xg for 20 min. The supernatants were dilute 10 times with 2% HNO3 (analytical grade) before analysis. Analysis was performed using a ThermoFisher iCap RQ Quadrupole – ICPMS in Kinetic Energy Dispersion Sensitive mode. Helium and Argon were used as collision gas and carrier gas separately. Calibration curve was constructed using Agilent Intelliquant 68 multi-element standard No.1 (IQ-1, #5190-9422). Data processing was conducted by the software Qtegra. ICP-MS measurements were conducted in triplicate. Biological replicates, which represent separate protein expression and purification experiments, are identified in text.

## Results

### AlphaFold2 structural predictions of secreted rust proteins

We used a structural prediction approach to compare secreted proteins from five rust species: *P. graminis* f. sp. *tritici* (*Pgt*, causing wheat stem rust disease); *P. striiformis* f. sp. *tritici* (*Pst*, wheat stripe rust); *P. triticina* (*Pt*, wheat leaf rust); *P. coronata* f. sp. *avenae* (*Pca*, oat crown rust); and the more evolutionarily divergent *M. lini* (*Mli*, flax rust). We first identified a total of 27,090 predicted secreted proteins from the genome annotations of the five species (*Pgt*: 6,891; *Pt*: 6,172; *Pca*: 6,018; *Pst*: 4,690; *Mli*: 3,319, Table S3), with 52.8% and 14.7% of these secreted proteins predicted as cytoplasmic and apoplastic effectors, respectively, by EffectorP 3.0 (Sperschneider & Dodds, 2022) (Table S4, S5). Approximately one in five of the secreted proteins (*Pgt*: 16.3%; *Pt*: 21.7%; *Pca*: 20%; *Pst*: 18.7%; *Mli*: 29.8%) are predicted to have PFAM domains. As expected, secreted proteins without PFAM domains are enriched for predicted effectors (cytoplasmic/apoplastic effectors: *Pgt*: 62.3%/16.8%; *Pt*: 59.3%/12%; *Pca*: 59.3%/12.6%; *Pst*: 58.8%/18.2%; *Mli*: 50.8%/20.1%) compared to those with PFAM domains (cytoplasmic/apoplastic effectors: *Pgt*: 30.8%/13.3%; *Pt*: 29.7%/10.7%; *Pca*: 27.6%/10.7%; *Pst*: 29.6%/13.2%; *Mli*: 22.9%/12.1%).

We predicted structures for these 27,090 secreted proteins using AlphaFold2, with 45.7% giving rise to predicted structures of high confidence (average pLDDT score >= 70), 35.2% medium confidence predictions (pLDDT score of 50 to 70) and 19% showing poor quality predictions (pLDDT score < 50) (Table S5). The distribution of pLDDT scores was broadly similar for the secretomes of each species (Figure S1). However, we observed significantly higher pLDDT scores for secreted proteins with PFAM domains than those without, both for predicted effectors and non-effectors (Figure 1, Figure S2). On average, the set of predicted non-effectors without PFAM domains had the lowest structure prediction scores.

**Figure 1:**
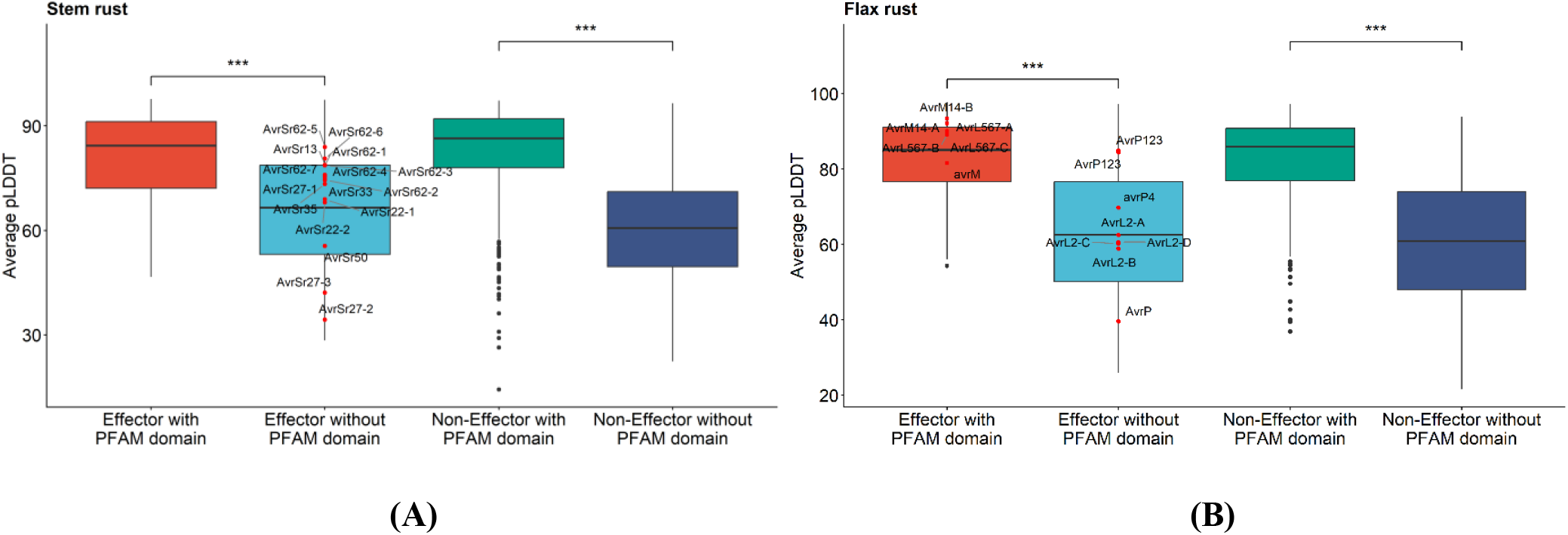
Confidence scores of AlphaFold2 predictions of (A) *Pgt* (stem rust) and (B) *Mli* (flax rust) secretomes. The box plots show the distribution of the average predicted local distance difference test (pLDDT) scores. Scores of described Avr effectors from the studied rust pathogens and their allelic variants are highlighted as red dots. The bounds of the box represent 25th to 75th percentiles, with the bold line highlighting the median. The whiskers are drawn to the smallest and largest data point that is no further than 1.5 * IQR from the hinge. Outliers are indicated by black dots.

Of 17 known *Pgt* effector proteins (AvrSr13, AvrSr22, AvrSr27, AvrSr33, AvrSr35, AvrSr50, AvrSr62 and their allelic variants), 12 had predicted structures of high confidence (Figure 1A). For *Mli*, 8 of the 14 known effector proteins had predicted structures of high confidence, while the AvrP4 prediction was of medium confidence and poor quality predictions were obtained for AvrP and AvrL2 alleles (Figure 1B). Seven effectors from *Pgt* and *Mli* have experimentally determined structures (AvrSr27-1, AvrSr50, AvrSr35, AvrP, AvrL567-A/D, AvrM-A/avrM, AvrM14-A/B) and as expected for the majority of these AlphaFold2 returns a structure of high confidence. However, the *Mli* effector AvrP has a poor prediction (pLDDT score 39.6) and the *Pgt* effector AvrSr50 has a medium confidence prediction (pLDDT score 55.5). Despite an experimentally determined structure and high confidence prediction for AvrSr27-1 (pLDDT score 74.6), the allelic variants AvrSr27-2 and AvrSr27-3 display poor predictions (pLDDT scores 34.4 and 42.1, respectively).

### Structural clustering reveals conserved and lineage-specific folds in rust fungal secretomes

To investigate structural conservation and diversity, we applied a structure-based clustering approach to the rust secretomes. In addition to the 27,090 AlphaFold2-predicted structures, we included 58 experimentally determined effector protein structures from various fungal and oomycete pathogens (Table S1), along with extracted structures of the individual N- and C-terminal domains of AvrSr27 and Avr1/SIX4, which each contain two subdomains. These included representatives from known structural families such as MAX, RALPH, FOLD, WY, ToxA-like and LARS. Because the rust genome references used here include two haploid nuclear genomes, many genes are expected to be represented by two allelic copies in each species and thus we defined a structural cluster as any group containing more than two proteins. Across the combined dataset of predicted and experimentally determined structures, we identified 860 structural clusters containing a total of 15,005 proteins (Table 1). There were also 1,682 doubletons, 97% of which are species-specific and likely represent allelic pairs. One-third of the combined secretome (*n* = 8,783) consisted of singletons that did not cluster with any other protein. Both singleton and doubleton proteins had significantly lower pLDDT scores than clustered proteins (Figure 2A), so their lack of clustering may result from inaccurate structural predictions. Many of the structural clusters were represented in the secretomes of multiple rust species (Figure 2B), including 67 clusters containing proteins from all five rust species and representing 6,205 proteins, comprising 41.4% of the 15,005 clustered proteins. Another 36 clusters containing 2,959 proteins (19.7% of clustered proteins) were represented in all four *Puccinia* species but not in flax rust. Each of the species also contained several species-specific structural clusters, although these contained relatively few proteins on average (Figure 2B).

**Table 1:** Structural cluster properties for the collective rust secretomes and 62 experimentally determined effector protein structures from various fungal and oomycete pathogens.

| Category | Number of structural clusters | Number of secreted proteins | % of the secretome |
| --- | --- | --- | --- |
| Protein singletons | 8,783 | 8,783 | 32.3% |
| Proteins doubletons | 1,682 | 3,364 | 12.4% |
| Proteins in clusters of 3+ | 860 | 15,005 | 55.3% |

**Figure 2:**
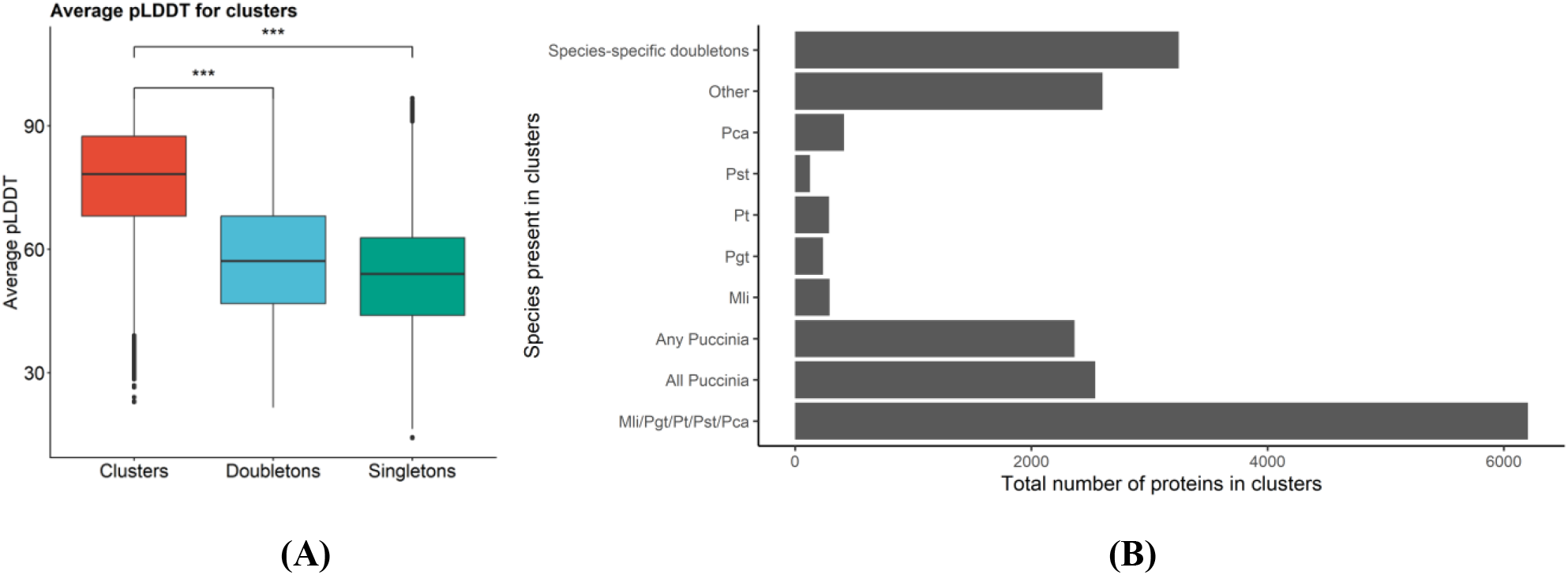
AlphaFold2 structural protein clusters of the collective rust secretomes. **(A)** The average predicted local distance difference test (pLDDT) scores for the AlphaFold2 predicted proteins for structural clusters (containing three or more family members), doubletons and singleton proteins. The bounds of the box represent 25th to 75th percentiles, with the bold line highlighting the median. The whiskers are drawn to the smallest and largest data point that is no further than 1.5 * IQR from the hinge. **(B)** Number of secreted proteins in structural clusters with different distribution patterns across the rust species, as indicated on the *y*-axis. ‘Other’ includes clusters present in *M. lini* and one to three of the *Puccinia* species.

### Some known effectors occur in large structural families

We observed 59 large structural clusters containing over 50 proteins each, with a maximum of 530 proteins in cluster #1 (Figure 3A, Figure S3). Fifteen of these clusters contain members with predicted PFAM domain functional annotations, such as clusters 3, 20, 29 and 36 which include predicted glycosyl hydrolases and clusters 25, 34 and 44 containing peptidases. Five of these large structural clusters contain known effectors, including the AlphaFold2 structure of AvrSr13 (cluster #7) (Arndell et al., 2024), and the experimentally determined structures of AvrSr35 (cluster #10) (Förderer et al., 2022; Zhao et al., 2022), SIX8 (cluster #12) (Yu et al., 2024), AvrSr50 (cluster # 45) (Ortiz et al., 2022), and ToxA/Avr2 (cluster #47) (Di et al., 2017; Sarma et al., 2005) (Figure 3B). Although the AlphaFold2 predicted structure of AvrSr50 occurred in a singleton due to its poor prediction, 39 proteins from *Pgt*, 17 proteins from *Pst* and 9 proteins from *Pt* are predicted to adopt a structure related to the experimentally determined structure of AvrSr50 (Figure 3B). In contrast, the AlphaFold2 structure of AvrSr35 clusters with the experimentally determined structure as part of an expanded family of 211 proteins; 74 from *Pgt*, 74 from *Pst*, 58 from *Pt* and 5 from *Pca* (Figure 3B). The AvrSr13 structural cluster #7 has expanded significantly in *Pgt* with 253 proteins, and includes the recently identified AvrSr33 (Spanner et al., 2026) as well as 22 proteins from *Pt*. The AvrSr50, AvrSr35 and AvrSr13 structural clusters are all absent from *Mli*.

**Figure 3:**
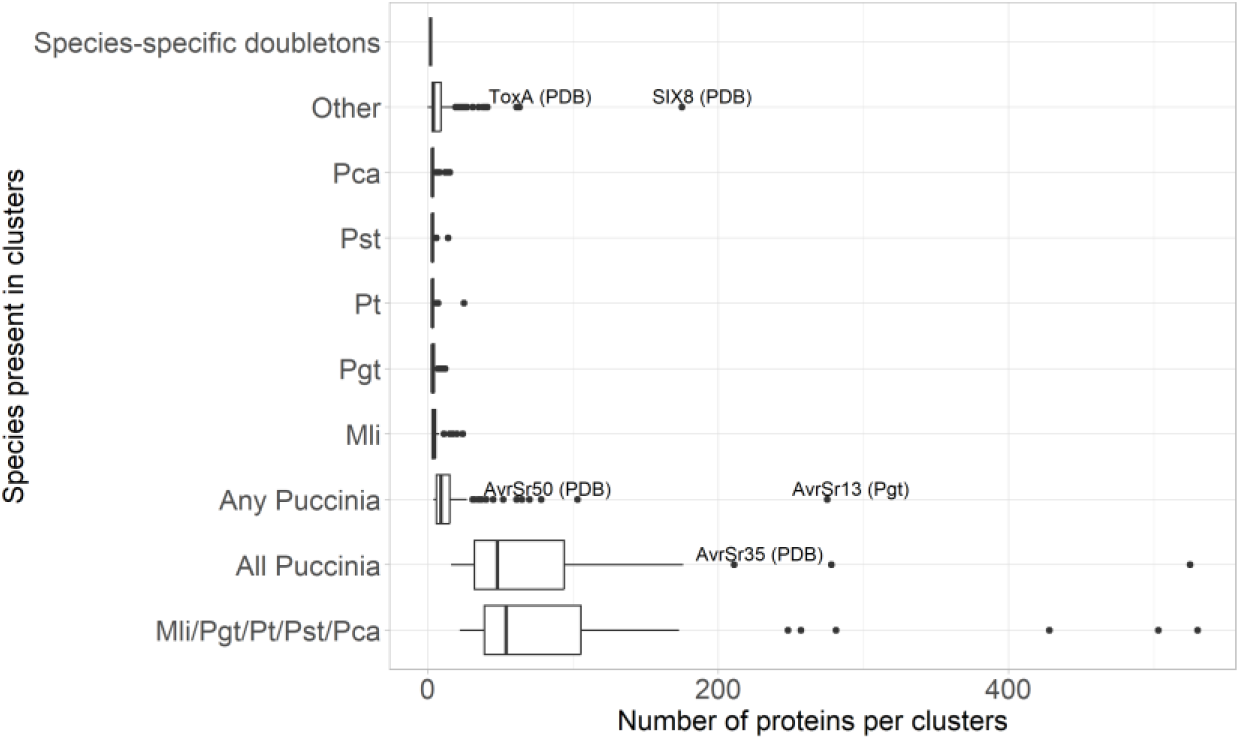
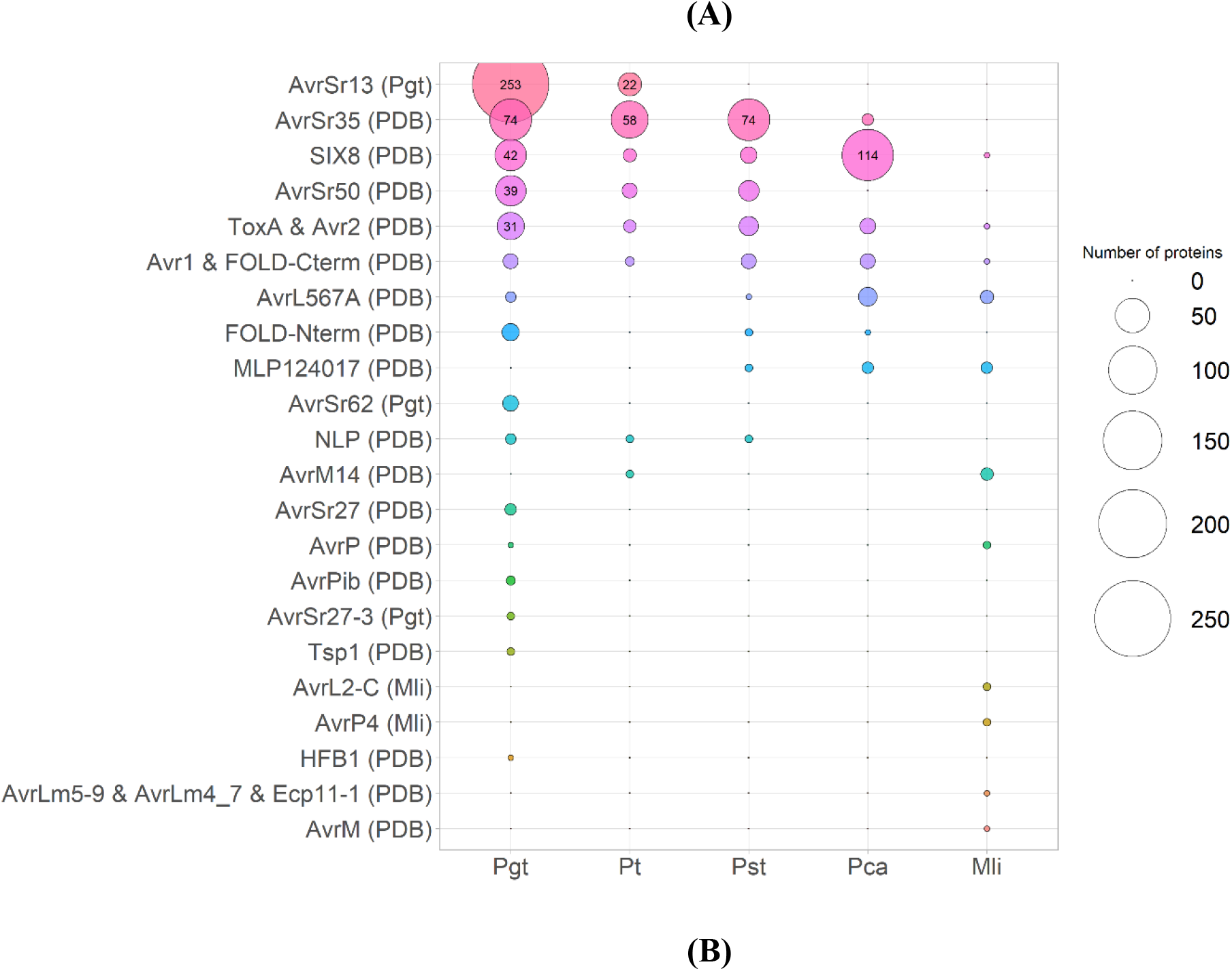
Distribution of structural clusters among rust secretomes. **(A)** The number of proteins per clusters and their conservations across species. Proteins with experimentally determined structures are represented by the label ‘PDB’. Some of the largest clusters contain known effectors (ToxA, SIX8, AvrSr50, AvrSr13, AvrSr35). **(B)** Representation of clusters containing known rust effectors or effectors from other plant pathogenic fungi or oomycetes across the five rust species. Circle represent the number of members of each cluster (*y*-axis) in the rust species (*x*-axis) according the scale at the right. Proteins with experimentally determined structures are represented by the label ‘PDB’.

Two other large clusters contain the experimentally determined structures of ToxA/Avr2 and SIX8. Although these have been suggested to be structurally related (Yu et al., 2024), here we found they occurred in two different clusters. SIX8 was part of cluster #12 with 114 proteins from *Pca* and 61 proteins from the other four rust species. The ToxA-like cluster #47 was smaller with 31 proteins from *Pgt* and 32 proteins across the other four rust species. The experimentally determined structure of AvrL567, which has also been suggested to be ToxA-like, despite some differences identified in the overall topology (Wang et al., 2007), forms a separate cluster with seven proteins from *Mli* (including the three AvrL567A-C proteins) and 19 proteins from *Pgt, Pst* and *Pca* (Figure 3B).

We also examined the clustering of the other experimentally determined or AlphaFold2-predicted structures of known effectors (Figure 3B). We did not detect any rust proteins in structural clusters containing the MAX, WY, RALPH, LARS or KP6-like effector structures, suggesting that these structural families are absent in these rust species. However, there were two clusters that share structural similarity to either the N- or C-terminal domains of the FOLD effector family (Figure 3B). The known *Mli* Avr effector proteins mostly belong to small clusters with few structural homologs, such as AvrM14 which is in a cluster of eight proteins including the experimentally determined structure. Despite sharing high sequence similarity, the four variants of AvrL2 (A-D) are structural singletons, possible due to their relatively poor AlphaFold2 prediction (Figure 1A, Figure S4). The AvrM AlphaFold2 structure formed a doubleton with the experimentally determined structure. The poorly predicted AvrP AlphaFold2 structure formed a singleton, whereas AvrP123 clustered with its experimentally determined structure and one other protein from *Pgt*. The two AvrP4 alleles formed a doubleton. The *Pgt* AvrSr62 effectors formed a cluster containing only the seven paralogous genes at the complex *AvrSr62* locus (Chen et al., 2025), while the two AvrSr22 alleles were singletons. The experimentally determined structure of AvrSr27 formed a cluster with five *Pgt* proteins but these did not contain the AlphaFold2 predicted structures of AvrSr27-1, AvrSr27-2 and AvrSr27-3. AvrSr27-2 formed a singleton whereas AvrSr27-1 and AvrSr27-3 formed a doubleton. Taken together, only the AvrSr50, AvrSr27-2 and - 3, AvrL2 and AvrP effectors did not cluster as expected with their experimentally determined structures and/or allelic variants, likely due to their low confidence predictions (Figure 1A, Figure S4).

### Rust pathogen secretomes contain expanded families of putative zinc-binding proteins

A high number of cysteine residues has been widely reported as a defining feature of effectors, with roles in disulfide bond formation shown in many apoplastic localising effectors (De Wit et al., 2009), or zinc coordination in a few cytoplasmic effectors (Outram et al., 2024; Zhang et al., 2018). About 73% of all proteins in the rust secretomes contain two or more cysteine residues (Figure 4A, Table S5), with predicted apoplastic effectors enriched for cys-containing proteins (88-95%) compared to those predicted to localise to the cytoplasm (60-67%; Figure 4B, Table S5).

**Figure 4:**
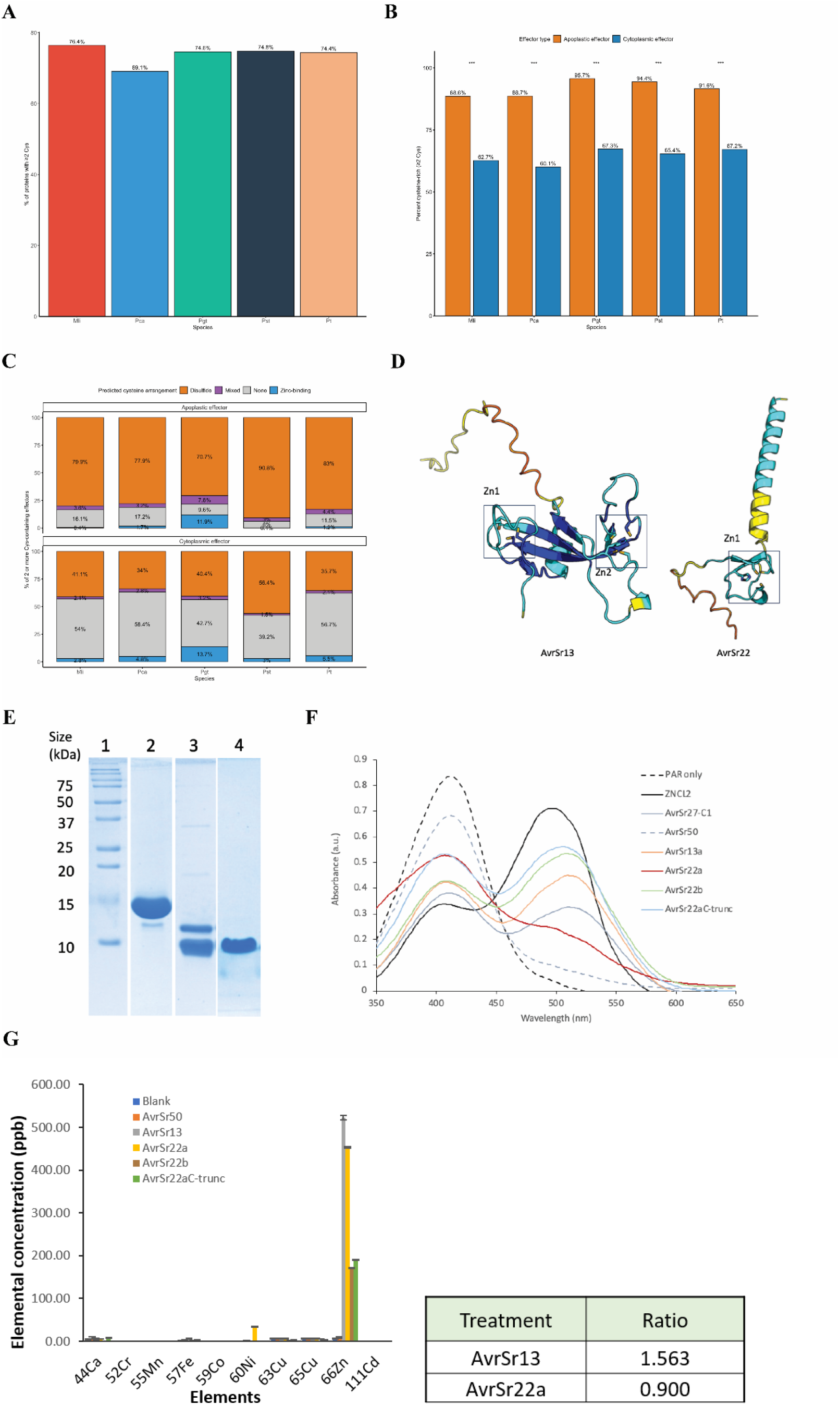
Cysteine-rich proteins in rust secretomes can be disulfide bonded or zinc bound. **(A)** Percentage of predicted effectors containing at least two cysteine across the five rust species. **(B)** Proportion of apoplastic versus cytoplasmic effectors within the cysteine-rich effector sets; significance assessed by χ^2^ test. **(C)** Predicted geometric arrangements of cysteine arrangements for apoplastic and cytoplasmic effectors that contain two or more cysteine residues: disulfide-bonded, zinc-binding, mixed (zinc + disulfide both identified) or none (no disulfide or zinc-binding identified. **(D)** Cartoon representations of the AlphaFold2 predictions of AvrSr13 and AvrSr22 coloured by per residue pLDDT score, where dark blue is the most confident, and yellow the least confident. Putative zinc binding pockets are highlighted by black boxes. **(E)** Coomassie-stained sodium dodecyl sulfate-polyacrylamide gel electrophoresis (SDS-PAGE) gel shows the purified proteins of AvrSr13, AvrSr22a and AvrSr22^C-trunc^ in lanes 2, 3 and 4 respectively. **(F)** Detection of metals in AvrSr13 and the variants of AvrSr22 using the 4-(2-pyridylazo) resorcinol (PAR). The absorbance spectra (350–650 nm) are shown for the following samples: PAR only, PAR plus ZnCl2, PAR plus AvrSr27-C1 (positive control), PAR plus AvrSr50-C (negative control), and PAR plus the test proteins AvrSr13a, AvrSr22a, AvrSr22b, and the AvrSr22a^C-trunc^, as indicated in the figure legend. **(G)** Quantification of metal ions (including Ca, Cr, Mn, Fe, Co, Ni, Cu, Zn, and Cd) in protein samples using ICP-MS. Protein samples (5 μM) were digested overnight at room temperature in 2% (v/v) HNO_3_ prior to analysis. “Blank” indicates the SEC buffer used for the protein. Error bars represent the standard deviation (SD) across three replicate injections. A summary table next to the graph presents the calculated zinc-to-protein molar ratios, offering a quantitative assessment of zinc binding per protein molecule.

Wehrspan et al. (2022) developed an *in silico* approach based on geometric arrangements of residues to detect zinc ion-coordination sites and disulfide bonds in predicted protein structures. We first used this method for the known zinc-binding effector proteins AvrSr27-1 and AvrP using their experimental structures (PDB ID: 8V1J and 5VJJ; modified to remove bound metals), and their AlphaFold2 predictions (including their alleles) (Table S6). This approach correctly identified the three and four zinc co-ordination sites in the AvrP and AvrSr27-1 experimentally determined structures, respectively (Figure S5A, Table S6). For the AvrSr27 predicted structures, only three of the four sites were predicted for AvrSr27-1 (Figure S5B, Table S6). Although the co-ordinating residues for Zn2 and Zn4 were correctly predicted, for the Zn1 site a different histidine residue (His33) was predicted to co-ordinate the zinc ion than observed in the experimental structure (His141) (Figure S6B). The Zn3 site was not predicted, likely due to the unusual side chain positioning of H68 in the AlphaFold2 model, which diverges from its position in the experimental structure (Figure S6A, B). The predicted structures of AvrSr27-2 and avrSr27-3 have lower pLDDT scores and only a single zinc co-ordination site was predicted for both, along with 1 or 3 disulfide bonds respectively (Figure S6B, Table S6). For the AvrP and AvrP123 AlphaFold2 structures, all three zinc binding sites were successfully identified (Figure S6C, Table S6) with no disulfide bonds. All five members within the AvrSr27-like cluster (#307) and the three proteins in the AvrP-like cluster (#585) were predicted to have at least one zinc binding site, though the number of sites predicted varied amongst the cluster members and some also contained predicted disulfide bonds (Table S5).

A total of 1,701 predicted protein structures across the five rust secretomes (∼6% of proteins) were predicted to contain at least one zinc-binding site, while 10,755 are predicted to contain disulfide bonds (Table S5), with 649 containing a mixture of both. Among effectors with two or more cysteine residues, apoplastic localising effectors were much more likely to contain disulfide bonds (70.7% to 90.8% in each rust species) than were cytoplasmic effectors (34% to 56.4% in each species; Figure 4C). The proportion of zinc-binding effectors was similar irrespective of localisation prediction (Figure 4C), indicating that zinc coordination is not strongly biased by predicted localisation. However, about three quarters of zinc-binding effectors are cytoplasmic, reflecting the larger number of cytoplasmic versus apoplastic effectors predicted in these secretomes. There was a notable enrichment of cys-rich proteins with neither disulfide bonds nor zinc binding sites in the cytoplasmic set (39.2%-58.4%) versus the apoplastic set (6.0%-17.2%). There was a substantially higher proportion of zinc binding proteins in *Pgt* than the other rusts (Figure 4D). This is largely due to the expansion of the AvrSr13-like effector family in *Pgt* (Table S5, Figure 3A), which accounts for about one-third of zinc-binding proteins in this species. Of the 275 proteins in this cluster, 271 were predicted to bind to zinc, with 248 of these containing two predicted sites as predicted for AvrSr13. Two proteins contain four predicted sites and visual inspection of these structures revealed they contained a domain duplication (Figure S6A-C) leading to the increased number of predicted zinc binding sites. AvrSr22 was also predicted to co-ordinate one zinc ion (Figure 4D), while AvrP4 from *Mli* was predicted to be a disulfide-bonded protein (Figure S7).

### Validation of zinc-binding in AvrSr13 and AvrSr22

To validate the *in silico* prediction of zinc-binding for AvrSr13 and AvrSr22, we expressed these proteins in *E. coli* BL21 (DE3) in media supplemented with a trace metal solution (Outram et al., 2024; Studier, 2005). AvrSr13 was purified via affinity and size-exclusion chromatography and could be observed as a single band on a Coomassie stained SDS-PAGE at the molecular weight consistent with the proteins expected size of 13.5 kDa (Figure 4E). Intact mass spectrometry analysis confirmed the identity of the protein, and that the protein was purified with the cysteine residues in a reduced form (Figure S8A,B). AvrSr22 was purified as a partially truncated protein (Figure 4E and Figure S9A). We also expressed and purified the alternative allele, AvrSr22b, which differs by 9 amino acids (Arndell et al., 2024), and found it was also similarly processed (Figure S9B). The banding patterns were consistent with a C-terminal truncation as both proteins could be purified via an N-terminal histidine tag (Figure S9). Intact mass spectrometry analysis of the proteins returned two predominant species, with masses consistent with the full-length mature protein and a smaller version truncated at residue 74 (Figure S10) and including the N-terminal zinc-binding site. We therefore designed another construct to express this truncated fragment, designated AvrSr22a^C-trunc^, which could be purified as a single species as shown by SDS-PAGE (Figure 4E). Next, we tested if the AvrSr13, AvrSr22a, AvrSr22b and AvrSr22a^C-trunc^ proteins purified from *E. coli* were bound to metal ions. We utilised a 4-(2-pyridylazo) resorcinol (PAR) assay, whereby the purified proteins were denatured to release any coordinated metals which can then cause a shift in the PAR absorbance peak due to the formation of a PAR-metal complex. This is demonstrated by incubation of PAR with ZnCl_2_ control (free PAR ∼416 nm; PAR-metal complex ∼500 nm) (Figure 4F). We also observed a shift consistent with the formation of a metal-PAR complex after incubation with the positive control, AvrSr27-1, a known zinc-binding protein (Outram et al., 2024). In contrast, AvrSr50 does not bind metal ions (Ortiz et al., 2022) and did not show a shift in absorbance. AvrSr13 and all the variants of AvrSr22 produced an absorbance shift consistent with a metal-PAR complex, confirming that these proteins contain bound metal ions. To identify the bound metals in AvSr13 and AvrSr22, we used inductively coupled plasma mass spectroscopy (ICP-MS). Both AvrSr13 and AvrSr22 variants were found to be bound to zinc ions (Figure 4G). We determined a metal ion occupancy rate for AvrSr13 of 1.44±0.24 ions/per protein based on three independent purifications (Figure S11). These data suggest that the AvrSr13 protein has multiple zinc binding sites, which is consistent with the AlphaFold2 model that predicts 2 putative zinc binding sites (Figure 4D). We could not calculate an accurate occupancy rate for the full length AvrSr22 due to the presence of mixed protein species of different sizes. However calculations of the AvrSr22a^C-trunc^ form suggest an occupancy of 0.58 based on one protein purification. These data are consistent with the AlphaFold2 model prediction that suggests AvrSr22 contains a single zinc-binding site in the N-terminal region of the protein (Figure 4D). AlphaFold3 has the capacity to model the likelihood of ion and small ligand binding (Abramson et al., 2024). When we modelled the AvrSr13 structure with two zinc ions we observed a high ipTM (interface-predicted TM) score of 0.91, with the Zn ions placed in the predicted coordination sites, further supporting our findings that AvrSr13 is a zinc binding protein with two coordination sites (Figure S12). Similarly, structural modelling of AvrSr22 in AlphaFold3 with a single zinc atom included returned an ipTM score of 0.9, with the zinc ion bound in the predicted configuration (Figure S12).

## Discussion

Effector proteins are key determinants of plant disease outcomes and are under evolutionary pressures, leading to high levels of sequence diversity. Experimental structure determination and computational predictions have suggested that many sequence-unrelated effectors belong to conserved structural families (Lahfa et al., 2024; Mark & Sylvain, 2023; Outram et al., 2024; Seong & Krasileva, 2021; Seong & Krasileva, 2023). Here, we used AlphaFold2 to predict protein structures for the secretomes of five rust pathogens to provide insights into the structural repertoire of effector proteins. 81% of the 27,090 secreted proteins gave rise to medium to high confidence structural predictions, which included most of the known Avr proteins identified in *Pgt* and *Mli*. However, the remaining 19% yielded low-confidence structural models, limiting their utility for detecting structural similarities. The accuracy of AlphaFold2 predictions is strongly influenced by the depth of evolutionary information available in multiple sequence alignments, and the presence of similar structures in the Protein Data Bank (Jumper et al., 2021). Effector proteins are known to be highly sequence diverse and typically yield poor multiple sequence alignments with few sequences present. Conversely, secreted proteins with functional annotations exhibited significantly higher structure confidence scores than those without, likely due to the increased depth of sequence alignment. However, it is also possible that some low-confidence models result from inaccurately annotated gene models or from long disordered regions. For example, the bacterial effectors AvrPto and AvrRpt2 have been shown by NMR spectroscopy to possess intrinsic disorder, with AvrPto containing a three-helix bundle core with long-disordered N- and C-termini (Wulf et al., 2004). Intrinsic disorder in effectors may enable flexible conformations to be adopted to facilitate interactions with multiple host targets.

Despite these limitations, a substantial portion of rust secreted proteins formed conserved structural clusters, offering a robust foundation for comparative and functional analyses across species. The experimentally determined structures of seven rust Avr effectors are all structurally distinct (Förderer et al., 2022; Ortiz et al., 2022; Outram et al., 2024; Zhang et al., 2018). In the present study, these effectors along with another six rust effectors all form discrete structural families, except for AvrSr13 and AvrSr33 which were both members of a single large cluster. This reinforces the previous finding that rust effectors are structurally diverse and contrasts with the findings in some other filamentous plant pathogens, such as *Blumeria* spp. where almost all known effectors fall into the RALPH family (Bauer et al., 2021) and oomycetes whose effector repertoire is dominated by WY class RxLR effectors (Boutemy et al., 2011). However, some rust effector structural families are greatly expanded at least in some rust species. For instance, AvrSr13 and AvrSr33 belong to a cluster containing 253 proteins in stem rust and 22 proteins in leaf rust, with no representatives in any of the other rust species. The AvrSr35-like family (211 proteins) occurs across all of the *Puccinia* species, but not in *Mli*. Of known structural families defined in other fungi, only the ToxA-like and FOLD families were detected in the rust secretomes. Interestingly, separate clusters were identified containing structures related to either the N-terminal or C-terminal domains of the FOLD family, suggesting these domains need not always be coupled. Although some previous reports suggested that many secreted proteins from *Pgt* and *Pt* contained Y/F/WxC motifs typical of powdery mildew RALPH effectors (Godfrey et al., 2010), we did not detect any rust proteins structurally-related to the RALPH family. Similarly, no representatives of the LARS, MAX, KP6, LysM or WY families were identified in these rust species.

Previously, we showed that two cytoplasmically localised cysteine-rich effectors (AvrP and AvrSr27) co-ordinate zinc ions through conserved cysteine and histidine residues (Outram et al., 2024; Zhang et al., 2018). Similar results were also reported for AvrPii from *M. oryzae* (De la Concepcion et al., 2024). This contrasts with the prevalence of disulfide bonds in cysteine-rich apoplastic effectors in other fungi (De Wit et al., 2009). Using an *in silico* analysis, we found that AlphaFold2 frequently identified the correct spatial arrangement of cysteines capable of forming zinc-binding pockets, even though the relevant ligands were not present in the input, and this could be distinguished from arrangements likely involving disulfide bonds. Cysteine-rich proteins predicted to have an apoplastic location overwhelmingly favoured spatial arrangements consistent with disulfide bonds (∼85%), while those with a cytoplasmic localisation were much less likely to contain disulfides (∼45%). Conversely, most zinc-binding effectors were predicted as cytoplasmic. Zinc-binding effectors appear to be overrepresented in the *Pgt* secretome, which partially reflects the major expansion of the AvrSr13/AvrSr33 family in this species (253 members). We confirmed the predicted zinc-binding status of two effectors, AvrSr13 and AvrSr22, by biochemical analysis *in vitro*. The involvement of four coordinating residues in AvrSr13 and AvrSr22, like that of AvrSr27 and AvrP (Outram et al 2024), suggests a structural rather than enzymatic role for the bound zinc ion(s), as the latter usually involve a labile fourth ligand or five co-ordination sites (Ataie et al., 2008; Mechtinger et al., 2025). However, the biological function and when these proteins acquire zinc remains to be understood. During preparation of this manuscript, AlphaFold3 was released, offering reported improvements in prediction of protein–protein and protein–ligand interfaces. Benchmarking AlphaFold3 on AvrSr13 and AvrSr22 showed high confidence scores for zinc ion coordination in the expected binding sites, suggesting that it could be a reliable tool for prediction of zinc-binding effectors. Overall, these observations highlight both the utility of structure prediction for identifying putative metal-binding or disulfide bonds in proteins and the need for experimental validation to resolve ambiguities. Protein structure predictions and relationships to known effectors may prove a useful additional metric for prioritising candidate effectors for downstream functional investigations.

## Author contributions

Conceptualization: JS, MF, MO, PND, SJW; Data curation: JS, MK, MO; Formal analysis: JS, MO, ZL; Funding Acquisition: JS, MF, PND, SJW; Investigation: JS, MO, ZL; Methodology:, JS, MK, MO, SJW; Project Administration: JS, MO, PND, SJW; Software: JS, MK; Resources: JS, MF; Supervision: JS, MF, MO, PND, SJW; Validation: JS, MF, PND; Visualization: JS, MO, ZL; Writing – original draft: JS, MF, MO, PND, SJW, ZL; Writing – review & editing: JS, MF, MO, PND, SJW, ZL.

