## Supplementary Figures for "Rust fungi secretomes contain structurally diverse effector families including cysteine-rich metal-binding proteins"

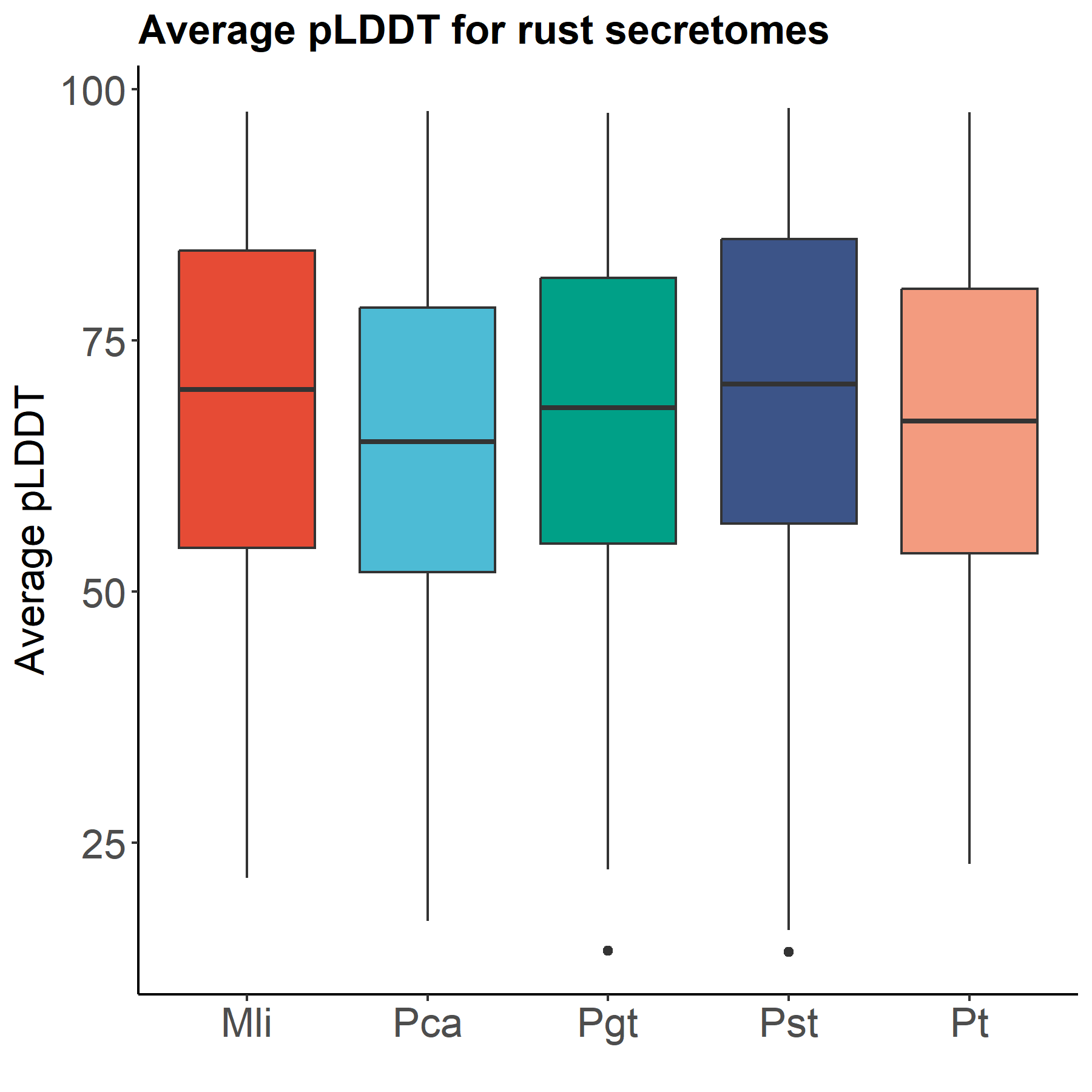
**Figure S1: Average pLDDT score distributions for the five rust secretomes.** The box plots show the distribution of the average predicted local distance difference test (pLDDT) scores for the predicted structures from each of the rust species, where an average score <50 indicates a low confidence prediction and >70 indicates a high confidence prediction. The bounds of the box represent 25th to 75th percentiles, with the bold line highlighting the median. The whiskers are drawn to the minima and maxima. Outliers are shown as black dots.

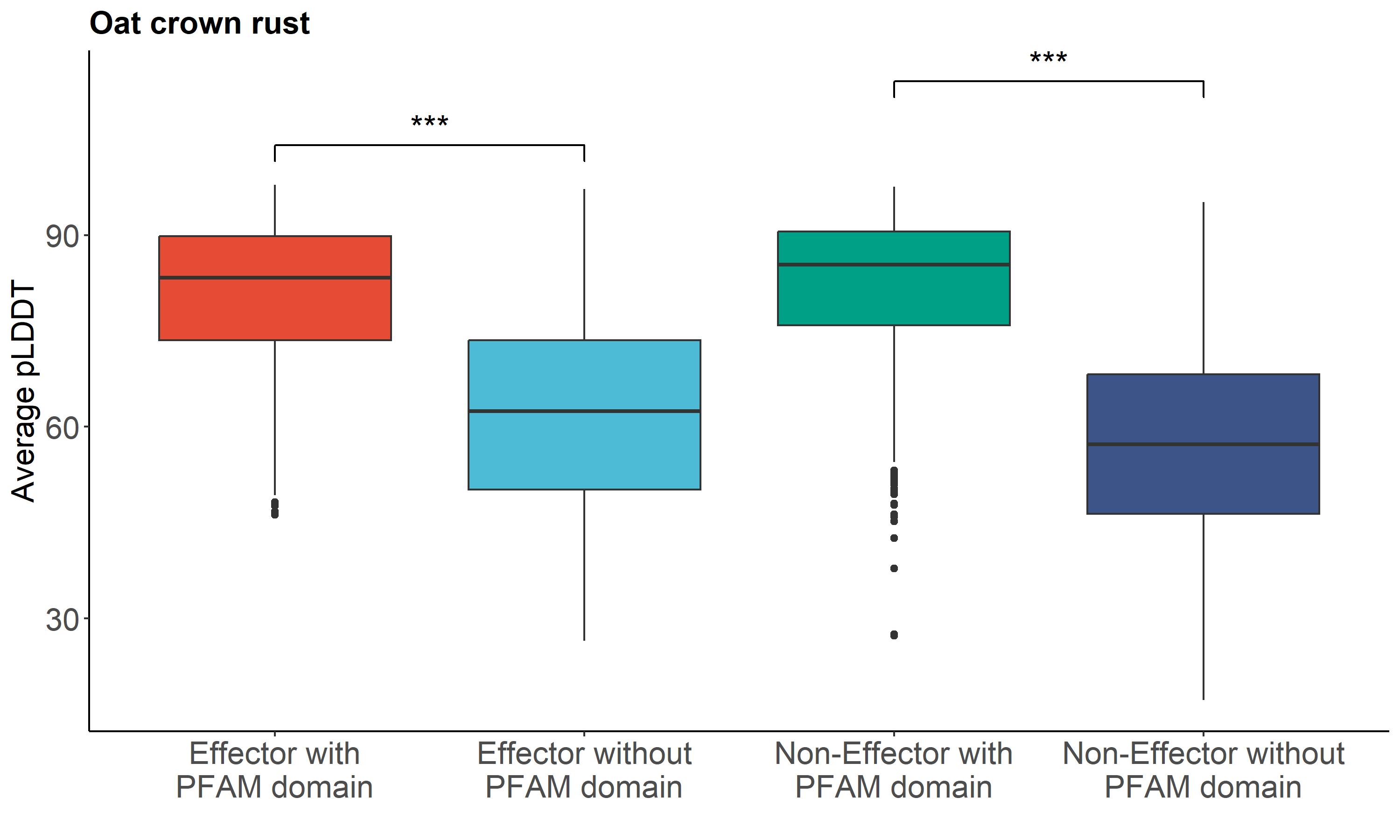

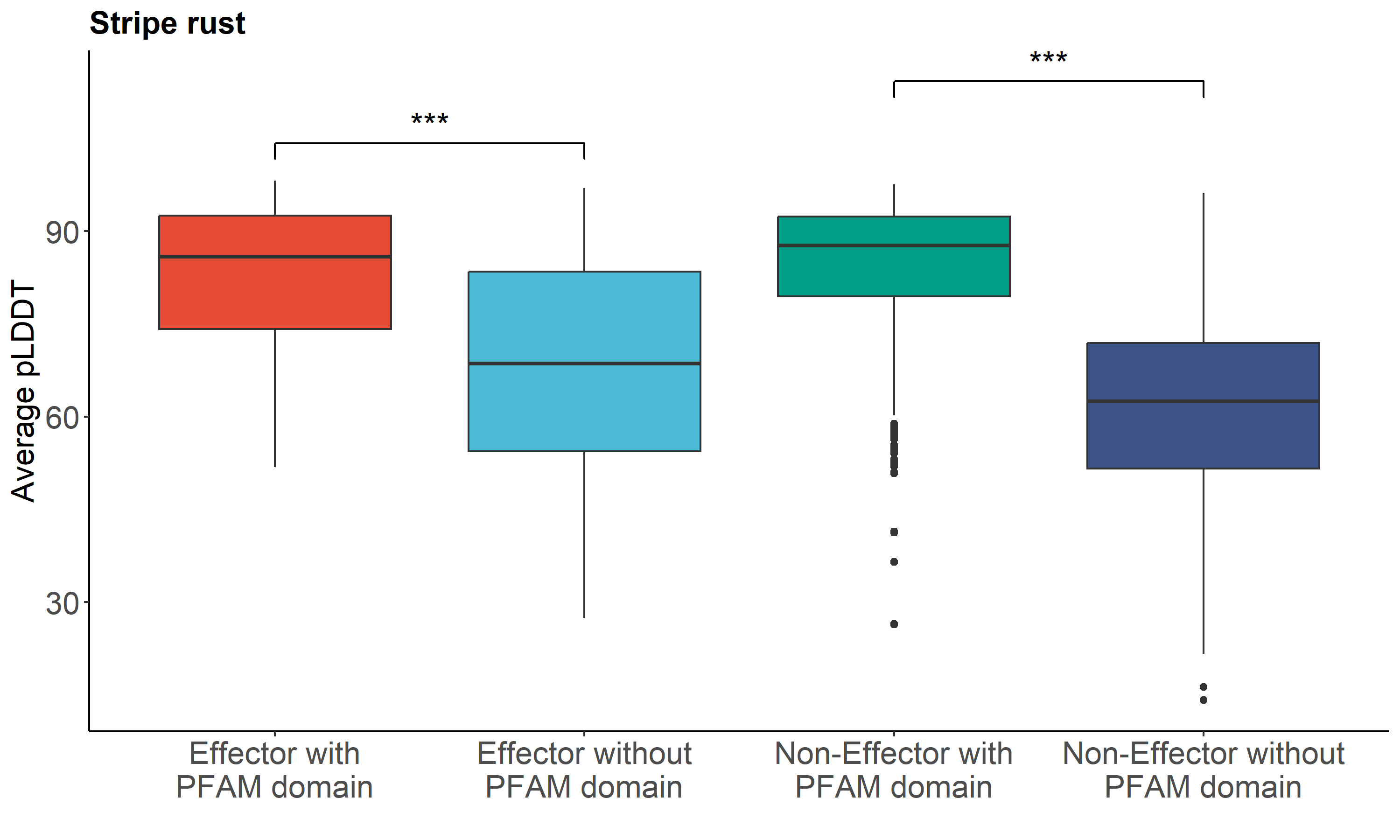

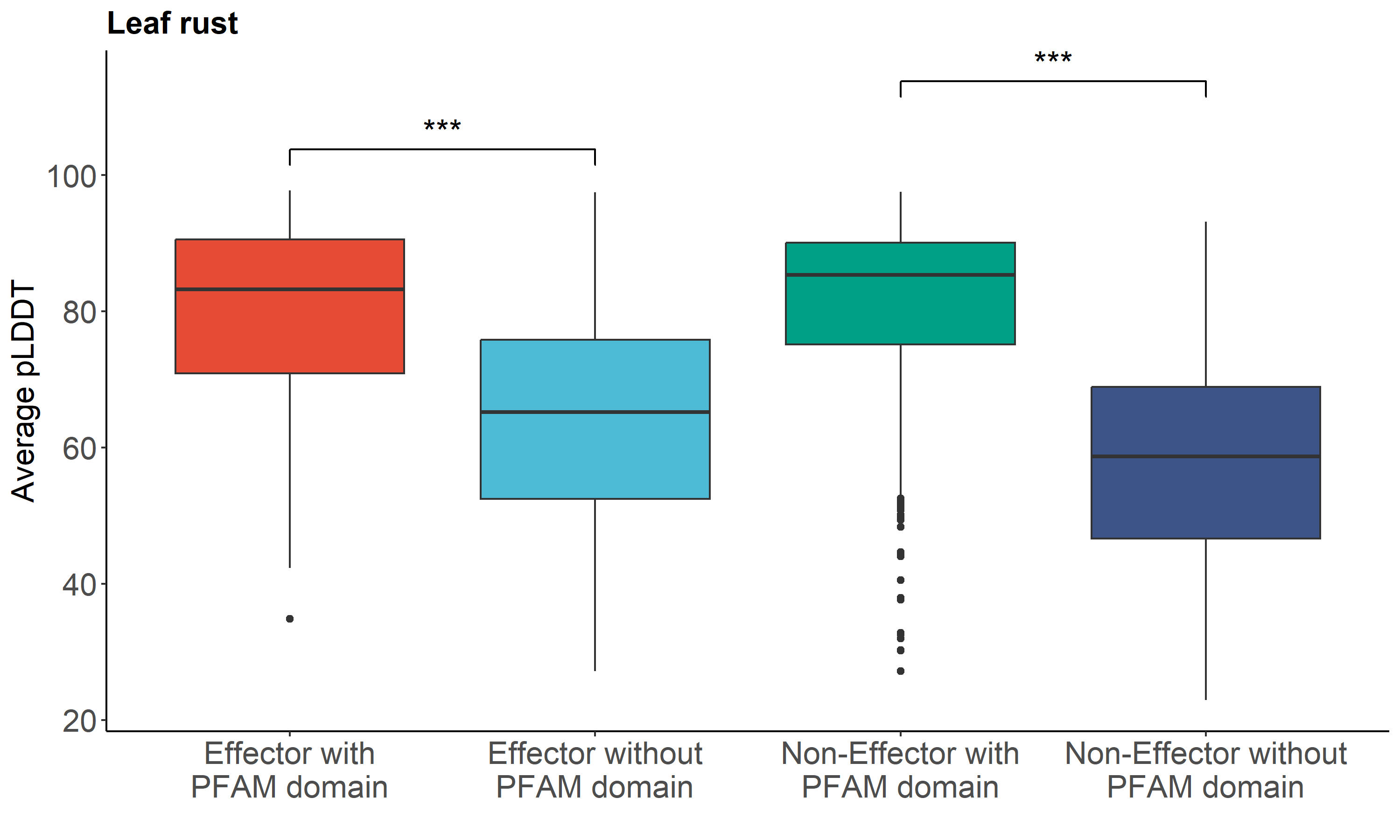

**Figure S2: AlphaFold2 predictions of *Pca* (oat crown rust), *Pst* (stripe rust) and *Pt* (leaf rust).** Secreted proteins with PFAM domains have significantly more confident AlphaFold2 predictions than those without PFAM domains. The box plots show the distribution of the average predicted local distance difference test (pLDDT) scores for the predicted structures from each of the rust species, where <50 indicates a low confidence prediction and >70 indicates a high confidence prediction. The bounds of the box represent 25th to 75th percentiles, with the bold line highlighting the median. The whiskers are drawn to the minima and maxima. Outliers are indicated as dots.

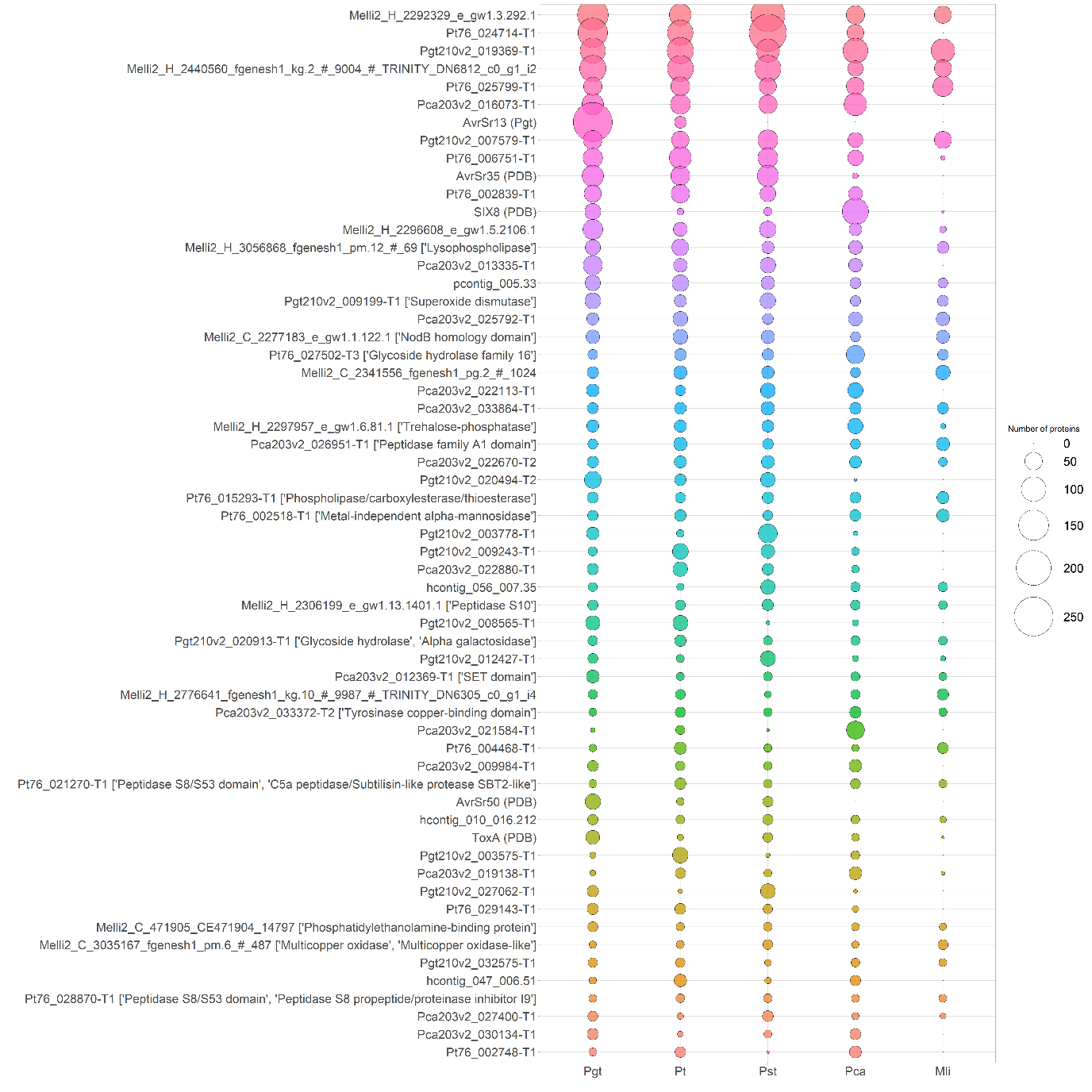

**Figure S3: Largest structural clusters identified in the rust secretomes.** Structural clusters with > 50 members are shown as a bubble plot. For each cluster, the corresponding identity of a representative protein is shown, alongside any known functions using sequence-based investigations.

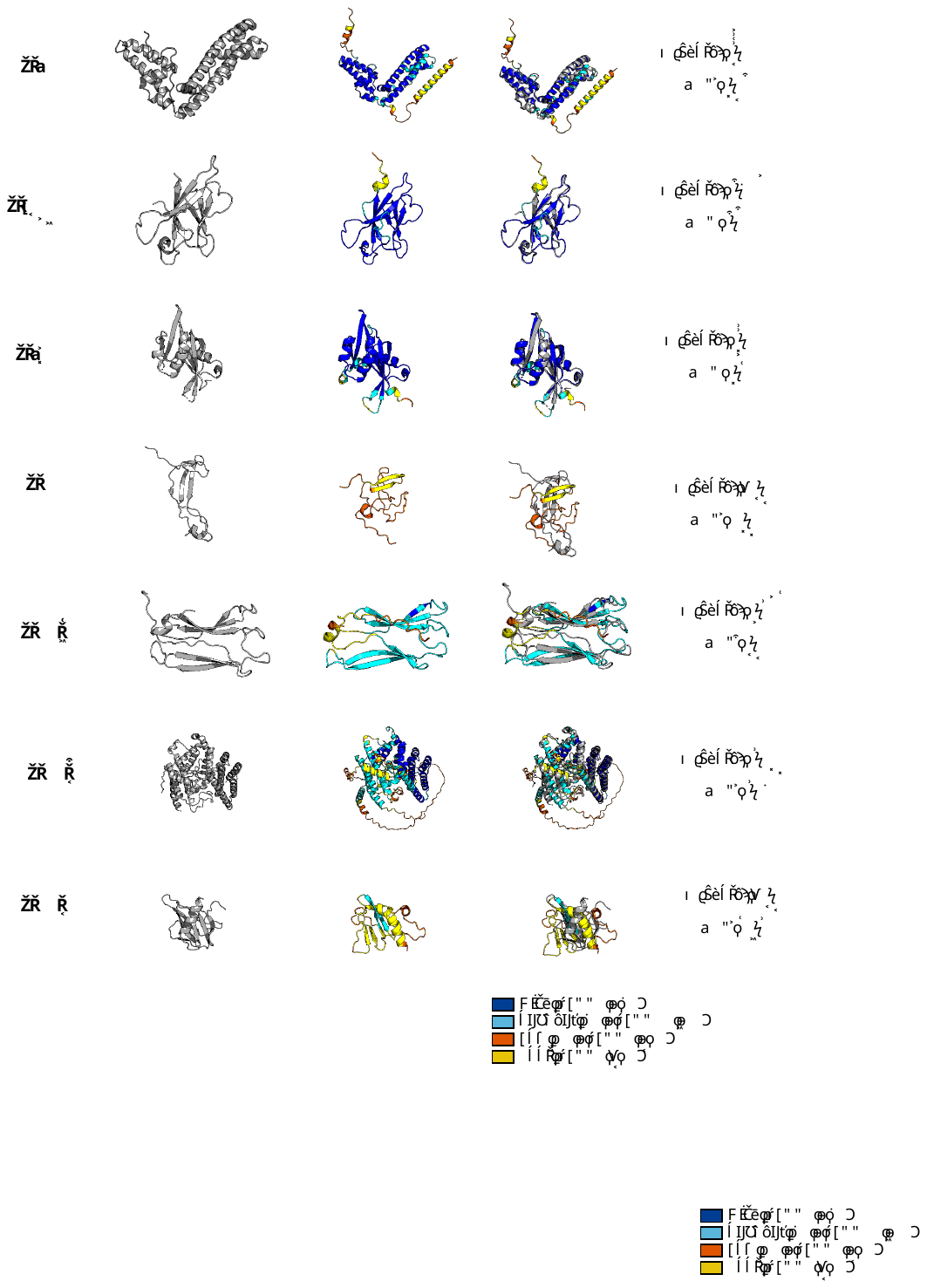

**Figure S4: Experimental structures of known Avr effectors. (A)** Experimentally determined structures for each Avr (left) shown in cartoon representation. **(B)** AlphaFold2 prediction of corresponding Avr, coloured according to average pLDDT score. **(C)** Overlay of experimental structure (grey) and AlphaFold2 prediction (coloured according to pLDDT score) with Tm-score and RMSD of alignment across the whole protein shown.

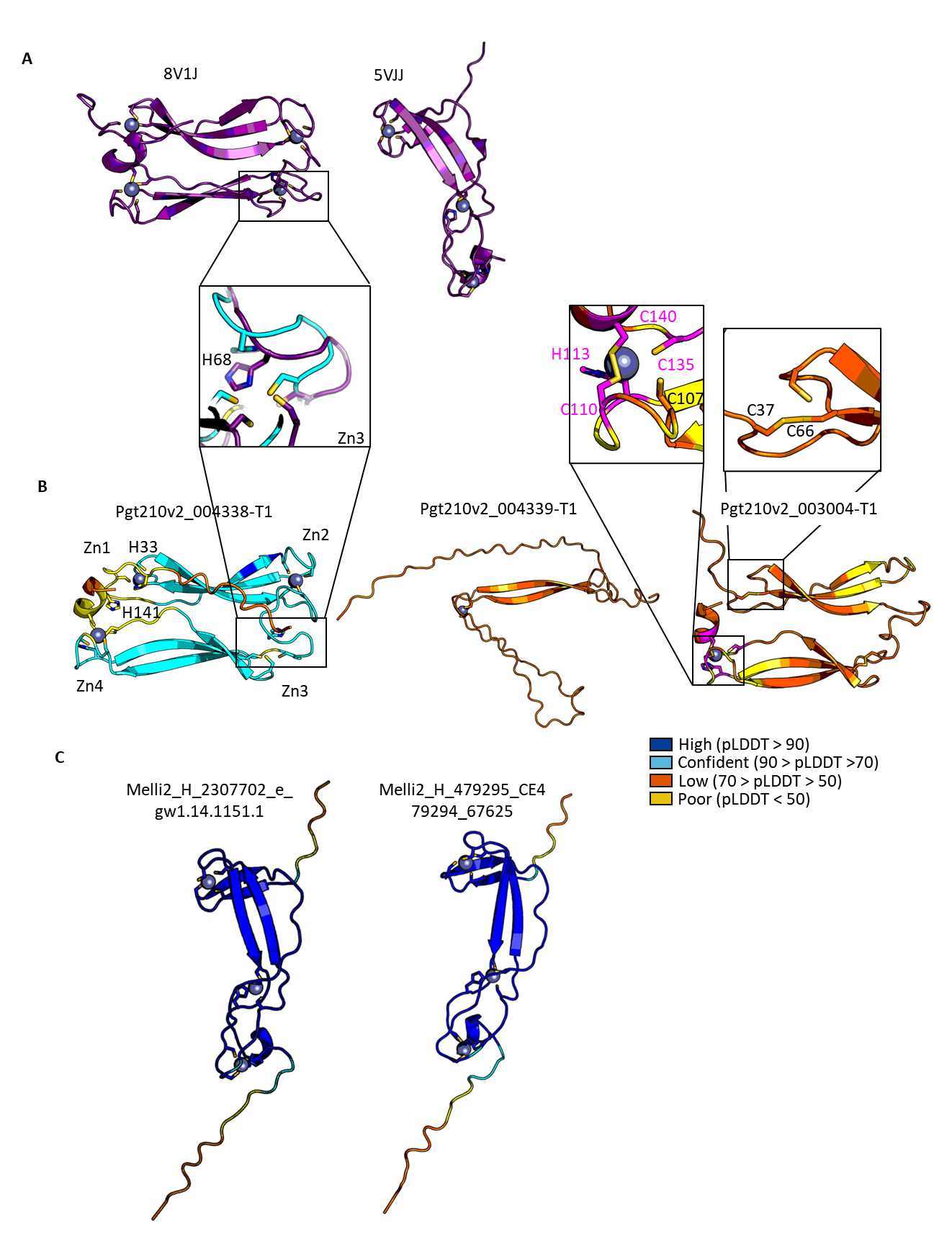

**Figure S5: Validation of the *in silico* metal binding prediction.** **(A)** Four zinc-binding sites were predicted for the crystal structures of AvrSr27 (8VIJ) and three zinc binding sites for AvrP (5VJJ). Both structures are shown in cartoon representation, with zinc molecules shown, and co-ordinating residues shown as sticks. **(B)** The AlphaFold2 predicted structures of the three AvrSr27 alleles shown in cartoon representation and coloured according to local confidence score (pLDDT), as represented in the legend. Predicted metal binding sites are shown with zinc ligand and co-ordinating residues. Insert panels show close up of zinc-binding site differences based on the experimental structure and the predicted model. **(C)** The AlphaFold2 predicted structures of AvrP-like effectors from the secretomes highlight that all three zinc pockets are predicted.

(A) **(B)**

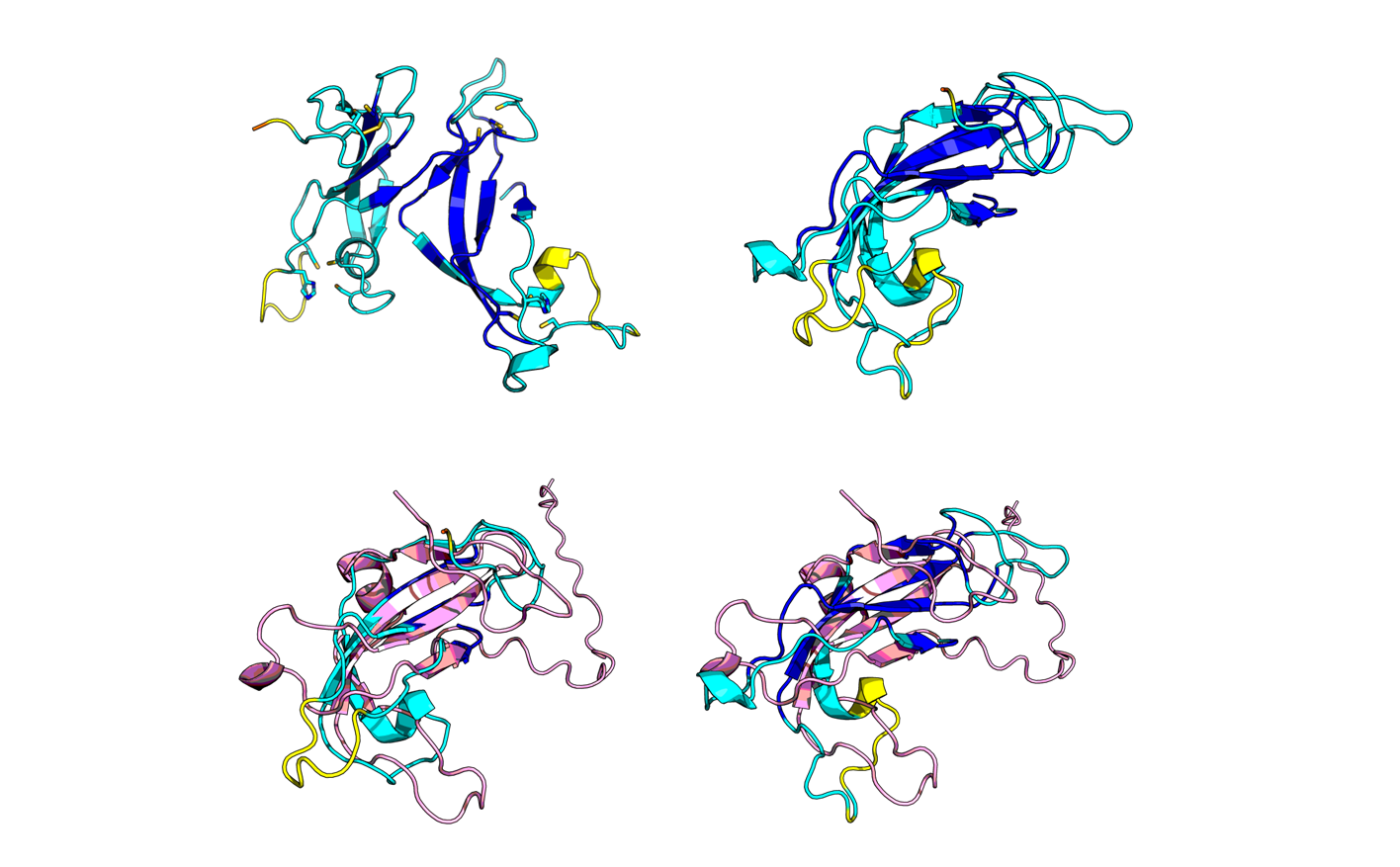

**(C)	(D)**

**Figure S6: AvrSr13-like effectors contain domain duplications. (A)** Cartoon representation of AvrSr13-like proteins identified to contain domain duplications of the “AvrSr13” domain, coloured according to pLDDT score. **(B)** Overlay of the two halves of the proteins from A, and with (**C)** AvrSr13 AlphaFold2 prediction shown in cartoon representation in pink showing structural conservation between the two domains and AvrSr13.

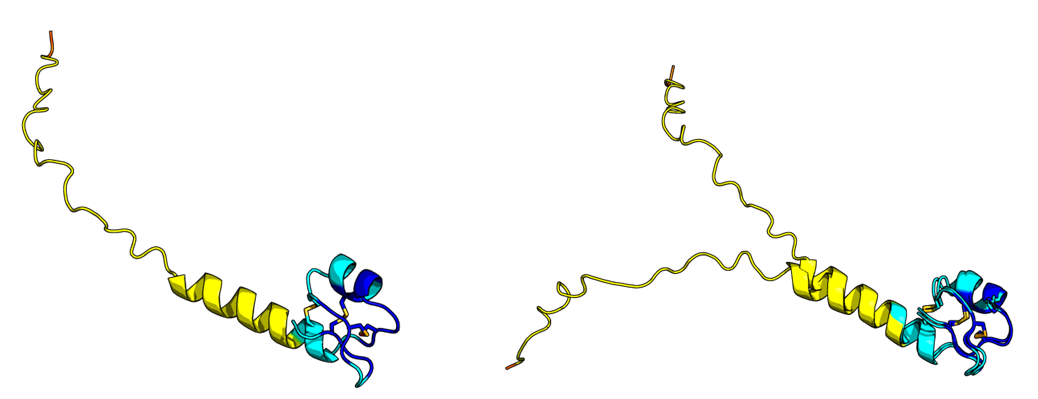

**Figure S7: AvrP4 is predicted by AlphaFold2 to be disulfide bonded.** A cartoon representation of the two AvrP4 alleles found in flax rust (Melli2_H_425135_CE425134_65539; Melli2_C_420024_CE420023_100808) coloured by pLDDT score. The predicted structure of AvrP4 has a disordered N-terminus, and six cysteine residues that are all predicted by AlphaFold2 to be involved in disulfide bond formation (shown as sticks coloured in yellow). This is in line with previous reports that suggest AvrP4 may form a cysteine knot based on cysteine residue spacing.

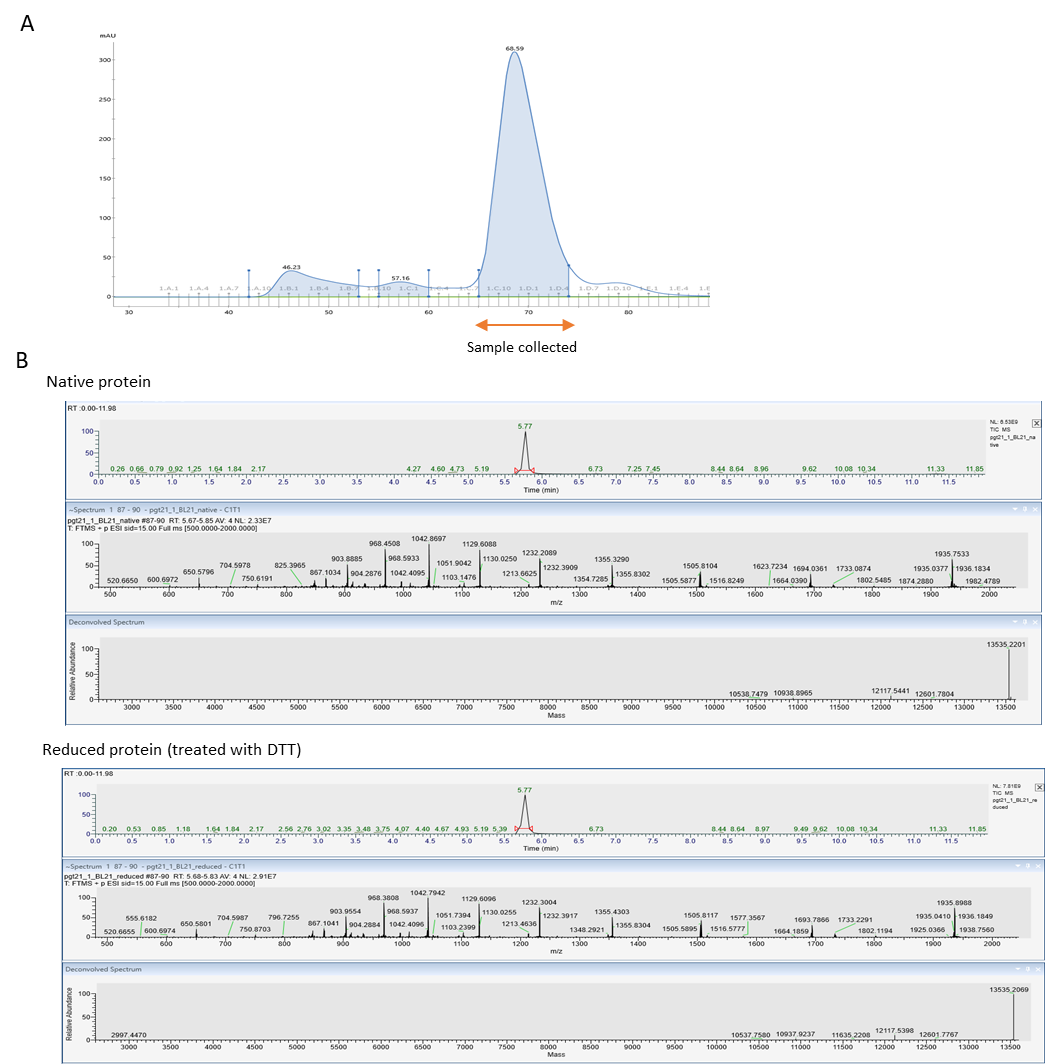

| **Protein** | **Theoretical molecular weight (Da)** | **Native protein** | **Reduced protein** |
| --- | --- | --- | --- |
| **AvrSr13a** | **13535.22** | **13535.22** | **13535.2** |

**Figure S8: SEC and intact mass spectrometry analysis of AvrSr13a protein.** **(A)** SEC chromatogram of AvrSr13a. The orange arrow indicates the elution peak corresponding to the pure protein fraction collected for further analysis. **(B)** Intact mass spectrometry of AvrSr13a under native and DTT-reduced conditions. The accompanying table compares the experimentally determined molecular weights with the theoretical values predicted by Peptide Mass.

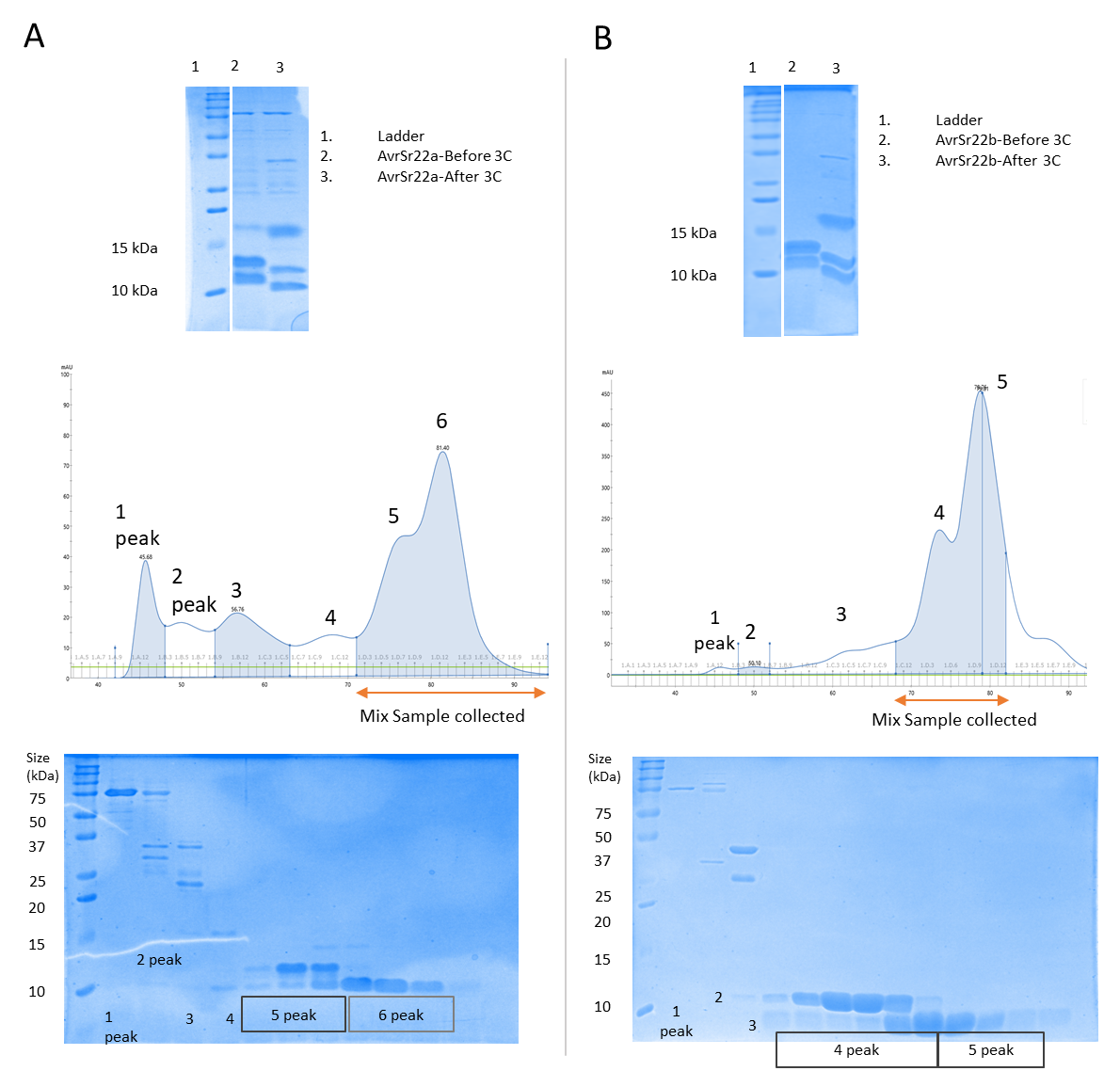

**Figure S9: Validation of AvrSr22 truncated.** **(A)** Analysis of AvrSr22a protein before and after 3C protease treatment. The top SDS-PAGE gel shows a shift in band position after treatment, indicating successful cleavage of the N-terminal tag. The middle panel shows the SEC profile of AvrSr22a, with all peaks labeled. The orange arrow indicates the fraction collected for downstream zinc-binding analysis. The bottom SDS-PAGE gel shows the protein content of the SEC fractions, demonstrating that full-length and truncated proteins could not be separated by SEC. **(B)** Same analyses performed for the AvrSr22b allele.

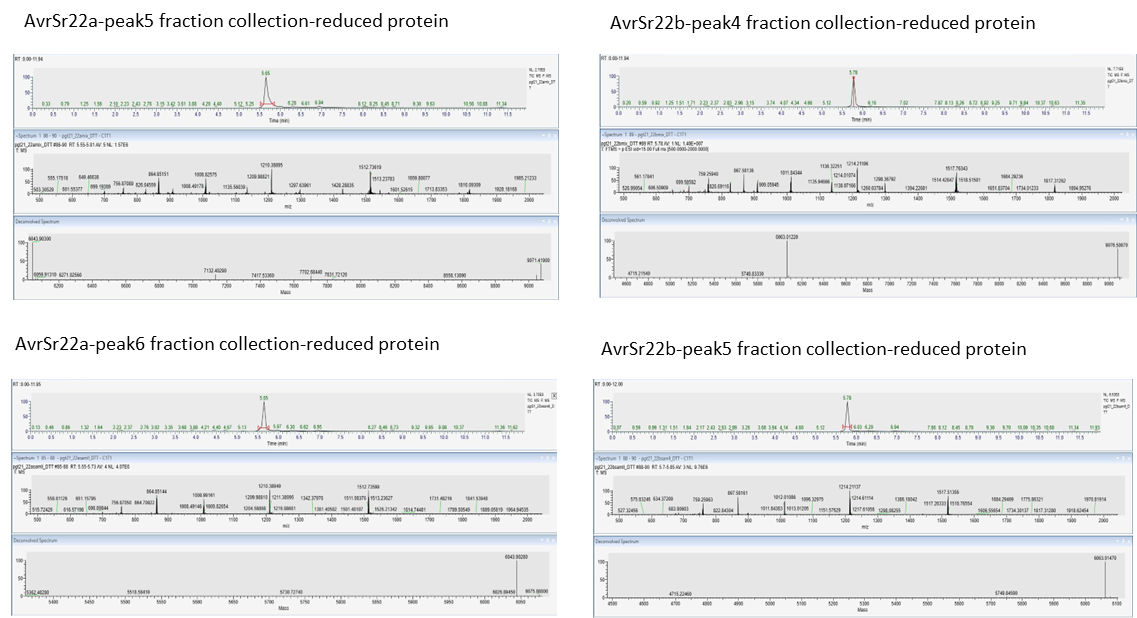

| **Protein** | **Theoretical molecular weight full length (Da)** | **Theoretical molecular weight for AvrSr22aC-trunc (Da)** | **Samples** | **Reduced protein** |
| --- | --- | --- | --- | --- |
| AvrSr22a | 9071.42 | 6043.91 | AvrSr22a-peak5 | 6043.9 |
|  |  |  |  | 9072.41 |
|  |  |  | AvrSr22a-peak6 | 6043.9 |
| AvrSr22b | 9076.51 | 6063.01 | AvrSr22b-peak4 | 6063.01 |
|  |  |  |  | 9076.5 |
|  |  |  | AvrSr22b-peak5 | 6063.01 |

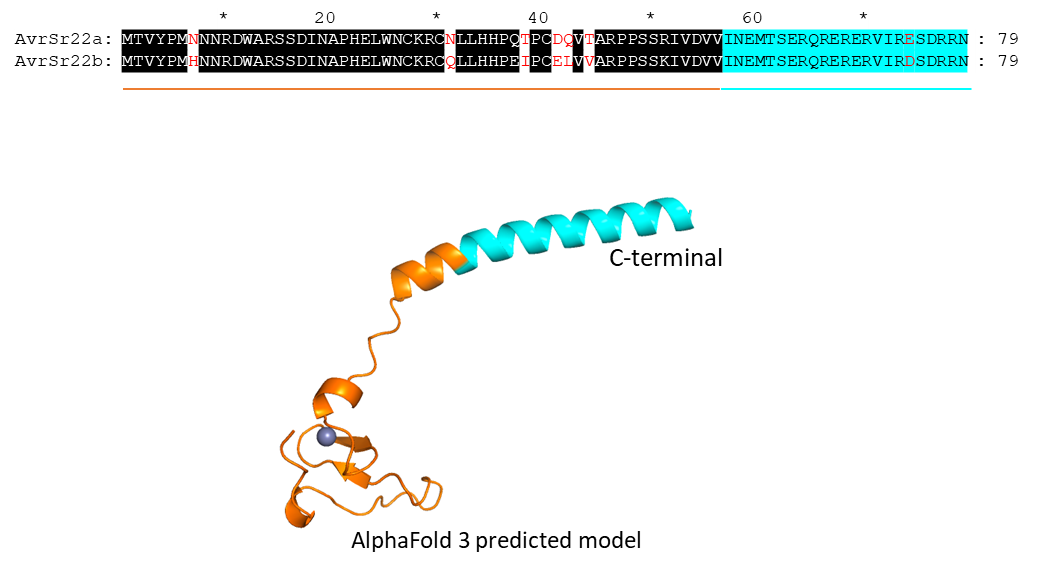

**Figure S10:** Intact mass spectrometry analysis of AvrSr22a (samples collected from SEC peak 5 and peak 6) and AvrSr22b (samples collected from SEC peak 4 and peak 5) under DTT-reducing conditions. The accompanying table compares the experimentally determined molecular weights with the theoretical values predicted by Peptide Mass. Amino acid sequence alignment of AvrSr22a and AvrSr22b is also shown, with the predicted cleavage sites indicated based on the observed molecular weights from intact MS. Both AvrSr22a and AvrSr22b consist of 79 amino acids, with the C-terminal 23 amino acids representing the truncated portion, highlighted in cyan in the alignment. An AlphaFold3-predicted structural model of AvrSr22b illustrates the location of the truncation in the 3D structure, with the truncated region colored in cyan to match the sequence alignment.

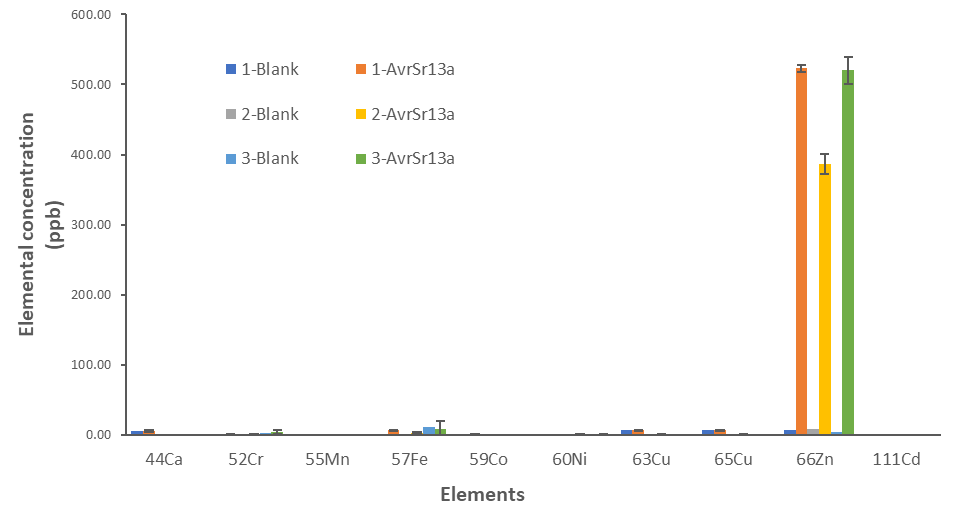

|  | **Ratio(Zn/proteins)** |
| --- | --- |
| 1-AvrSr13a | 1.58 |
| 2-AvrSr13a | 1.17 |
| 3-AvrSr13a | 1.58 |
| Average | 1.44 |

**Figure S11 Quantification of metal ions (including Ca, Cr, Mn, Fe, Co, Ni, Cu, Zn, and Cd) in AvrSr13a from three independent purifications, analyzed using ICP-MS.** “Blank” refers to the SEC buffer used for the protein. Error bars represent standard deviation (SD) from three replicate injections. The table shows the calculated zinc-to-protein occupancy.

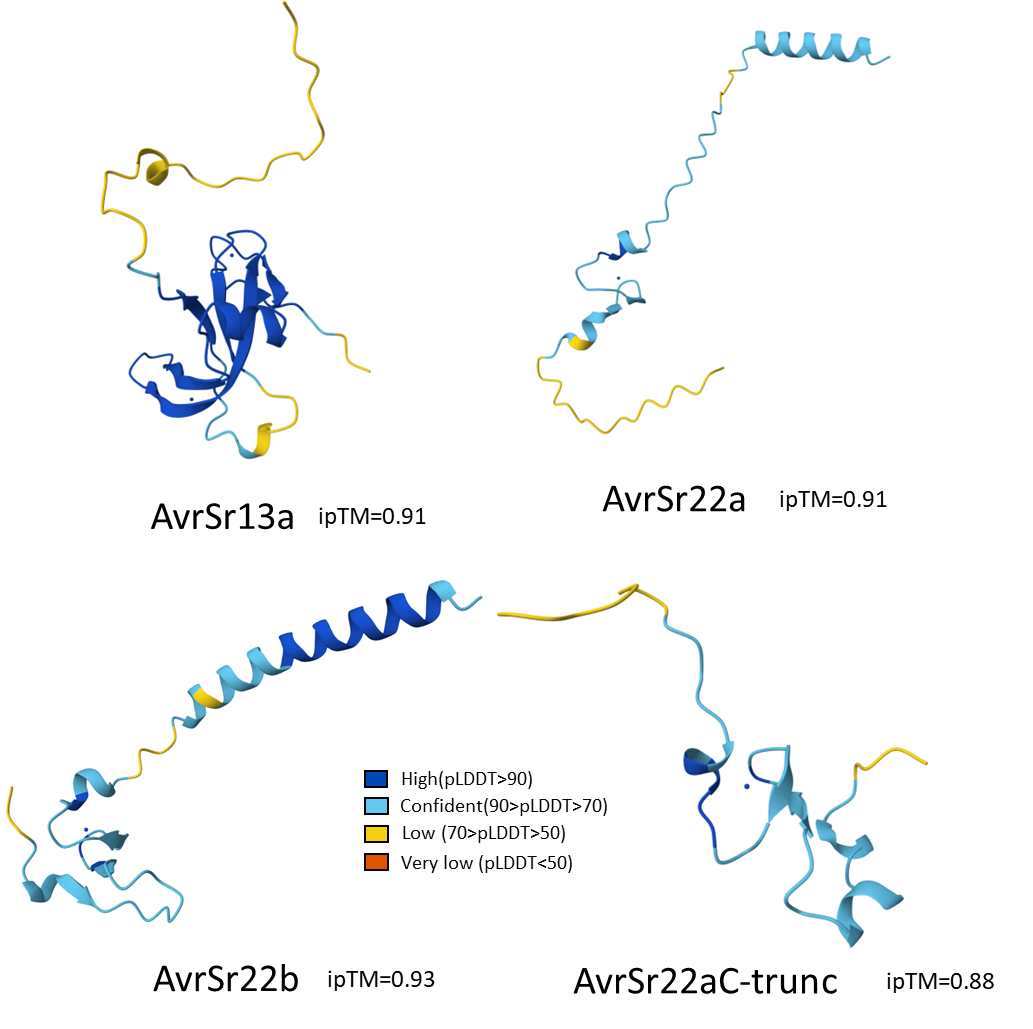

**Figure S12: AlphaFold3 predicted structures for AvrSr13a, AvrSr22a, AvrSr22b and AvrSr22aC-trunc.**
